# Mesophilic inmates of a geothermal vent-head as providers of critical ecosystem services

**DOI:** 10.64898/2026.05.19.726215

**Authors:** Subhajit Dutta, Aditya Peketi, Sumit Chatterjee, Jit Ghosh, Sai Pavan Pillutla, Nibendu Mondal, Mahamadul Mondal, Jagannath Sarkar, Swapneel Saha, Amlanjyoti Dhar, Aninda Mazumdar, Ranadhir Chakraborty, Wriddhiman Ghosh

## Abstract

Geothermal vent-heads exclusively support the proliferation of (hyper)thermophilic microorganisms. However, diverse mesophiles are stochastically thrust into these ecosystems by local geodynamic forces, and their population-level responses to the unfamiliar stressor hold critical implications for life’s intrinsic resilience to biophysical adversities. Here, we report a time-course exploration of the boiling vent-water of a Trans-Himalayan geothermal spring to determine the population dynamics, *in situ* functionalities, and potential ecological roles of mesophilic incomers in the context of indigenous (hyper)thermophiles. The temporally-consistent (hyper)thermophilic core-community, predominated in turn by a few Aquificia and Thermoproteota, was delineated as **m**etagenome-**a**ssembled **g**enomes. Mesophiles/moderate-thermophiles were identified as **MAG**s and/or pure-cultures, and whether temporally-consistent/inconsistent, had far lower prevalence. Overall, *in situ* transcriptional activity and prevalence had significant positive correlation; but many species exhibited high activity despite low prevalence and vice versa, implicating the decoupling of global metabolism from growth/proliferation. Bulked metatranscriptomic signatures identified the (hyper)thermophiles as primary producers of the ecosystem. Populations of mesophiles and moderate-thermophiles were apparently active, but not growing, *in situ*, with some being engaged in environment detoxification, and production of nutrients, metabolites and protective-biomolecules wanting in the (hyper)thermophiles. The findings engender an unprecedented portrayal of geothermal-vents as multi-community ecosystems where unfamiliar immigrants provide critical metabolic services.

## Introduction

High-temperature habitats are widely acknowledged as potential nurseries of early life on Earth^1,2^; yet, hot spring waters, compared to other aquatic ecosystems, harbor lower microbial diversities due to the biophysical constrains imposed by heat on the cell systems of a large majority of extant microorganisms^3–5^. The limited numbers of bacterial and archaeal taxa which encompass species inherently capable of growing at high temperatures, and so regarded as thermophilic, are thought to be the natural inhabitants of hot spring ecosystems^6–8^. That said, culture-independent explorations of geographically discrete geothermal ecosystems have revealed the presence of mesophilic bacteria not only in the spring water outflows^9–11^ and adjoining features^4,12–15^, but also within the boiling vent-waters discharged at the spring-heads^4,10,16–19^. A few instances of mesophilic bacteria being isolated as pure-cultures from boiling vent-waters or fumaroles^20–22^ are also there.

Population ecology of mesophilic microorganisms that are thrust into unfamiliar geothermal environments, often stochastically^23^ by local geodynamic forces^9–11^, holds critical implications for life’s intrinsic resilience to biophysical adversities. However, almost no information is available regarding the relative abundance and populational consistency of mesophilic microorganisms within geothermal-vent habitats. Moreover, there is complete dearth of knowledge on the *in situ* metabolic status and ecological fate of the immigrants that inadvertently find their way into these unfamiliar extreme environments.

Here we temporally resolve the population dynamics of the mesophilic (and moderately-thermophilic) microorganisms present at the vent-head (discharge epicenter) of a Trans-Himalayan geothermal spring, in the quantitative context of the hyperthermophilic and thermophilic species dwelling in the same habitat. The species-level microbiome architecture of the boiling vent-water of the explored spring - situated in the **C**humathang **g**eothermal **a**rea (**CGA**) of Eastern Ladakh at an altitude of 3950 m above the mean sea-level^17,24^ - was revealed alongside its chemistry over six sampling occasions, spanning a period of one year. Over the first four investigations, a general consistency was discovered in the chemical characteristics of the vent-water, alongside an overall uniformity in the identity of the **m**etagenome-**a**ssembled **g**enomes (**MAG**s) retrieved. Consequently, through the fifth and sixth rounds of investigation, mesophilic bacteria were isolated and characterized genomically, in tandem with which *in situ* functionalities of all the cultured and uncultured species discovered were determined via metatranscriptome analysis.

The results illustrate that the temperature-affinity-based microbial communities of a geothermal vent have highly asymmetric (disproportionate) prevalence and functionality, not only between them, but also within the species assemblages. Whereas the preponderant (hyper)thermophilic community accomplishes primary production (chemolithotrophic energy capturing and autotrophic carbon fixation) in the ecosystem, the mesophilic minority remains active sans growth, and renders secondary metabolic services crucial for the ecological efficacy of the hyperthermophiles. The findings show wild populations of mesophilic bacteria to be far more heat-resilient than laboratory cultures, and in doing so modify our perception about geothermal-vent habitats from exclusive abodes of specialized extremophiles to more bio-diverse, inclusive, and interactive ecosystems. Adaptabilities of natural mesophilic populations to extreme biophysical conditions can help us design stress-resilient microbial systems for various biotechnological applications.

## Results and Discussion

### Study plan

Over separate forenoon and afternoon investigations carried out on 01-November-2021, 04-April-2022, and 01-November-2022 (Table 1; Fig. 1), boiling water from the vent epicenter of the most vigorous geyser of CGA situated on the bank of the river Indus (and referred to hereafter as the **C**humathang **m**ain **s**pring on **I**ndus or **CMSI**) was studied for its chemistry and microbiome architecture. This generated six independent sets of physicochemical data, microbial cell density records, and collections of MAGs or population-genome bins. Furthermore, through the forenoon and afternoon of 01-Nov-2022, the vent-water microbiome was investigated via bulked metatranscriptome analysis, and pure culture isolation and genome analysis.

**Figure 1.**
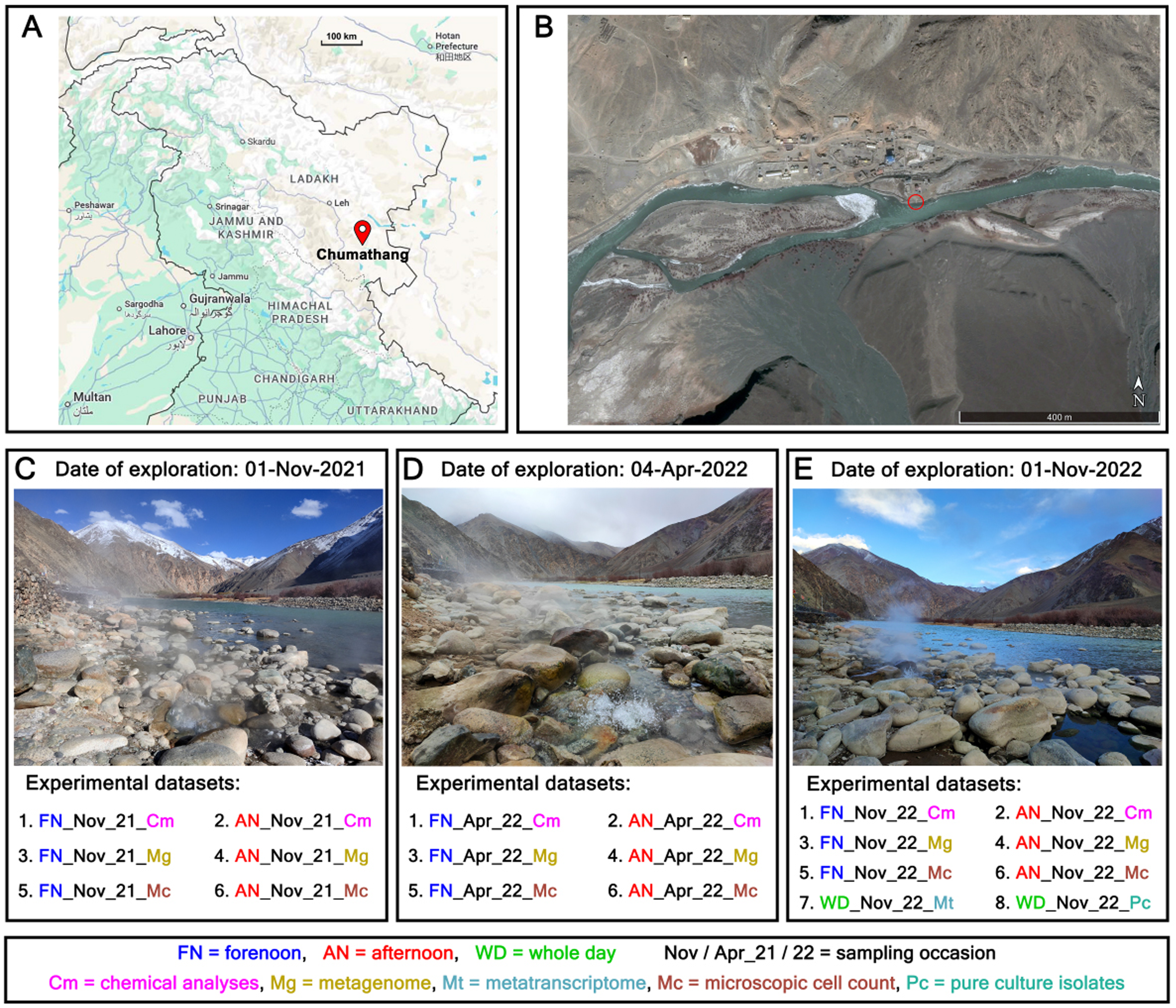
Geographical location and topography of the study site depicted in conjunction with a summary of the different types of data generated over the exploration. (**A**) Map of northern India (sourced from the publicly available database http://maps.google.com/) showing the position of the Chumathang area, within the territory of Ladakh. (**B**) Satellite image of the **C**humathang area (sourced from the publicly available database https://www.google.com/earth/) showing the position of the **m**ain **s**pring on river **I**ndus (CMSI). (**C** through **E**) Images of CMSI taken on the three different exploration days. Experimental information generated on each exploration day have been shown with dataset names indicating the hour of sampling (forenoon, afternoon, or whole day), occasion of sampling (November or April of 2021 or 2022) and the topic of investigation (chemistry, metagenomics, metatranscriptomics, microscopic cell count, or pure-culture isolation).

**Table 1.**
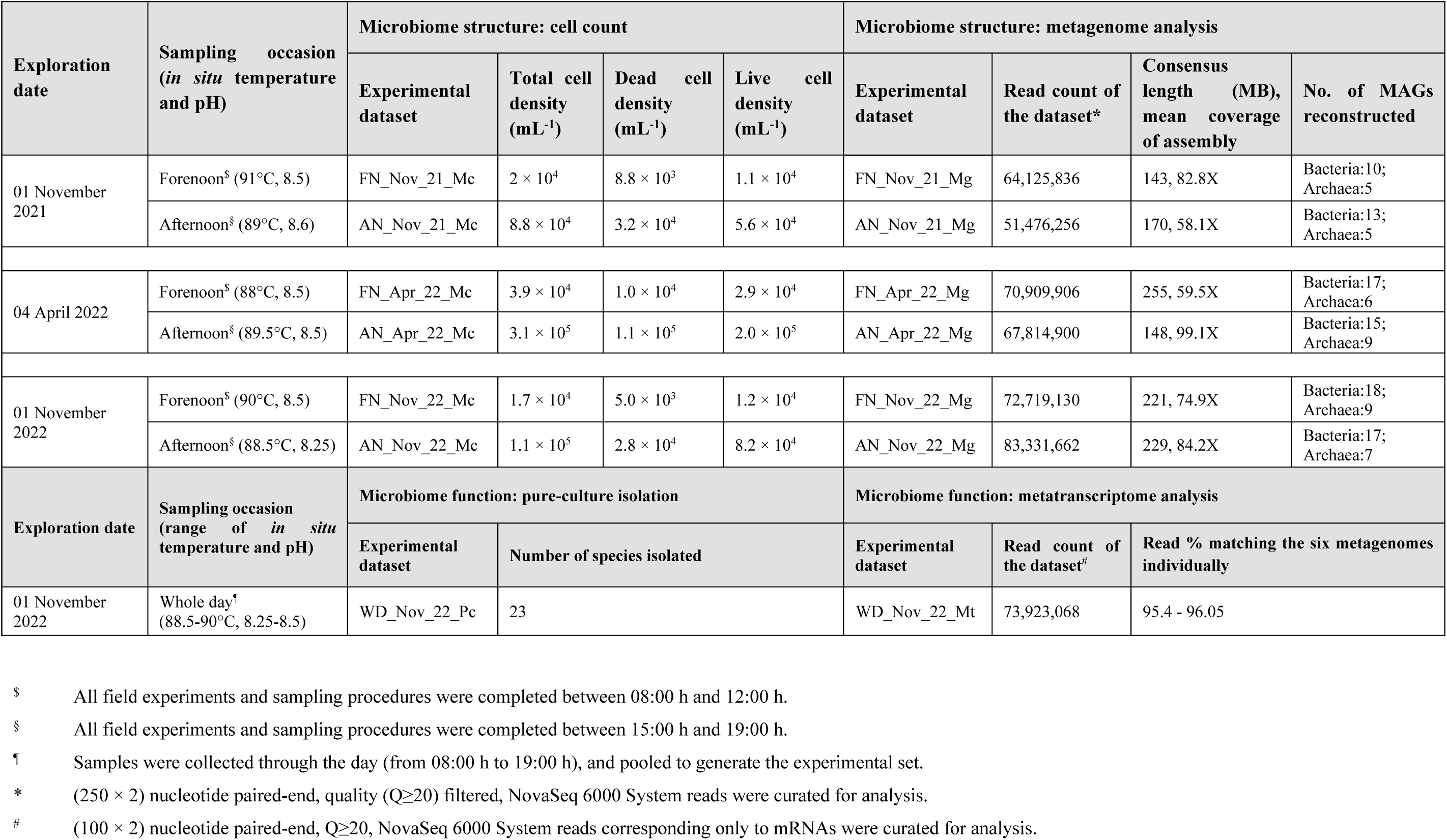
Summary of the geomicrobiological exploration carried out for the CMSI vent-water between November-2021 and November-2022.

| Exploration date | Sampling occasion<br>(*in situ* temperature and pH) | Microbiome structure: cell count |  |  |  | Microbiome structure: metagenome analysis |  |  |  |
| --- | --- | --- | --- | --- | --- | --- | --- | --- | --- |
|  |  | Experimental dataset | Total cell density (mL<sup>-1</sup>) | Dead cell density (mL<sup>-1</sup>) | Live cell density (mL<sup>-1</sup>) | Experimental dataset | Read count of the dataset* | Consensus length (MB), mean coverage of assembly | No. of MAGs reconstructed |
| 01 November 2021 | Forenoon<sup>§</sup> (91°C, 8.5) | FN_Nov_21_Mc | $2 \times 10^4$ | $8.8 \times 10^3$ | $1.1 \times 10^4$ | FN_Nov_21_Mg | 64,125,836 | 143, 82.8X | Bacteria:10; Archaea:5 |
| | Afternoon<sup>§</sup> (89°C, 8.6) | AN_Nov_21_Mc | $8.8 \times 10^4$ | $3.2 \times 10^4$ | $5.6 \times 10^4$ | AN_Nov_21_Mg | 51,476,256 | 170, 58.1X | Bacteria:13; Archaea:5 |
| 04 April 2022 | Forenoon<sup>§</sup> (88°C, 8.5) | FN_Apr_22_Mc | $3.9 \times 10^4$ | $1.0 \times 10^4$ | $2.9 \times 10^4$ | FN_Apr_22_Mg | 70,909,906 | 255, 59.5X | Bacteria:17; Archaea:6 |
| | Afternoon<sup>§</sup> (89.5°C, 8.5) | AN_Apr_22_Mc | $3.1 \times 10^5$ | $1.1 \times 10^5$ | $2.0 \times 10^5$ | AN_Apr_22_Mg | 67,814,900 | 148, 99.1X | Bacteria:15; Archaea:9 |
| 01 November 2022 | Forenoon<sup>§</sup> (90°C, 8.5) | FN_Nov_22_Mc | $1.7 \times 10^4$ | $5.0 \times 10^3$ | $1.2 \times 10^4$ | FN_Nov_22_Mg | 72,719,130 | 221, 74.9X | Bacteria:18; Archaea:9 |
| | Afternoon<sup>§</sup> (88.5°C, 8.25) | AN_Nov_22_Mc | $1.1 \times 10^5$ | $2.8 \times 10^4$ | $8.2 \times 10^4$ | AN_Nov_22_Mg | 83,331,662 | 229, 84.2X | Bacteria:17; Archaea:7 |
| Exploration date | Sampling occasion<br>(range of *in situ* temperature and pH) | Microbiome function: pure-culture isolation |  |  |  | Microbiome function: metatranscriptome analysis |  |  |  |
|  |  | Experimental dataset | Number of species isolated |  |  | Experimental dataset | Read count of the dataset<sup>#</sup> | Read % matching the six metagenomes individually |  |
| 01 November 2022 | Whole day<sup>¶</sup> (88.5-90°C, 8.25-8.5) | WD_Nov_22_Pc | 23 |  |  | WD_Nov_22_Mt | 73,923,068 | 95.4 - 96.05 |  |

<sup>§</sup> All field experiments and sampling procedures were completed between 08:00 h and 12:00 h.981 <sup>§</sup> All field experiments and sampling procedures were completed between 15:00 h and 19:00 h.982 <sup>¶</sup> Samples were collected through the day (from 08:00 h to 19:00 h), and pooled to generate the experimental set.983 <sup>\*</sup> (250 × 2) nucleotide paired-end, quality (Q≥20) filtered, NovaSeq 6000 System reads were curated for analysis.984 <sup>#</sup> (100 × 2) nucleotide paired-end, Q≥20, NovaSeq 6000 System reads corresponding only to mRNAs were curated for analysis.

### The low salinity, alkaline, but sulfate-rich, vent-water of CMSI features a stable physicochemical milieu

Across the six sampling occasions, temperature (88–91°C) and pH (8.25–8.6) of the fluid discharged at the vent-head showed little variation. Total alkalinity (7.5–7.7 mM), dissolved solids (1441–1674 ppm), salinity (757–839 ppm), and conductivity (1.64–1.75 mS) also exhibited only small levels of fluctuation (Table S1; Fig. 2A). Likewise, the most abundant solutes maintained more or less uniform abundances (Table S1; Fig. 2B) – of these, again, chloride and sodium concentrations fluctuated the least, followed by sulfate and silicon (no sulfide was detected in the vent-water on any sampling occasion); fluctuations in the concentrations of potassium and lithium were relatively higher, and those of boron and vanadium were the highest. These key physicochemical properties remained not only stable throughout the exploration but also within the ranges delineated previously for different CGA hot springs^17,25^. Salinity of the discharged fluid was low by global fresh-water standards^26^. However, its total alkalinity - theoretically attributable to high dissolved bicarbonate concentration acquired via subsurface geomorphic weathering - was far above the levels typically encountered for geothermal discharges^27^.

**Figure 2.**
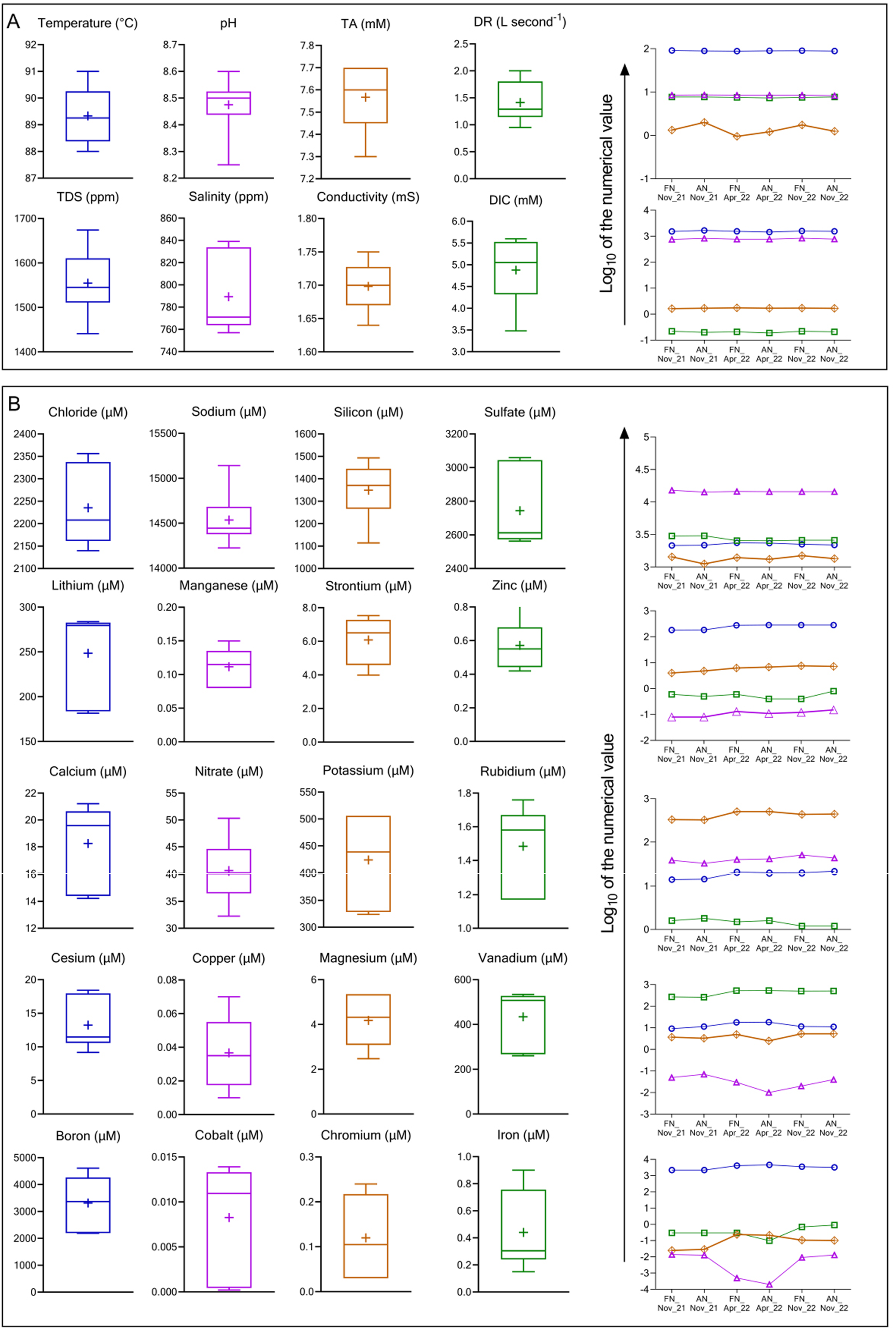
Physicochemical properties of the CMSI vent-water recorded over the six sampling occasions. (**A**) Box-and-whisker plots for temperature, pH, total alkalinity, discharge rate, total dissolved solids, salinity, conductivity, and dissolved inorganic carbon concentration of the water discharged at the epicenter of the CMSI vent. (**B**) Line plot representations for the parameters shown in panel “A”. (**C**) Box-and-whisker plots depicting the concentrations of chloride (Cl^-^), sodium (Na), silicon (Si), sulfate (SO_4_^2-^), lithium (Li), manganese (Mn), strontium (Sr), zinc (Zn), calcium (Ca), nitrate (NO^3-^), potassium (K), rubidium (Rb), cesium (Cs), copper (Cu), magnesium (Mg), vanadium (V), boron (B), cobalt (Co), chromium (Cr), and iron (Fe). (**D**) Line plot representations for the parameters shown in panel “C”. In panels “A” and “C”, each box represents the interquartile range capturing the central 50% of the data, with the horizontal line and the ‘‘+” symbol indicating the median and mean values respectively. The two whiskers extend up to the maximum and minimum values recorded for the parameter in question, above the third quartile and below the first quartile, respectively. In panels “B” and “D”, the six data points available for each parameter have been connected to enable easier tracking of the changes in the parameter; every parameter can be identified by its symbol as well as line color matching the color used in the corresponding box-and-whisker plot.

Notwithstanding the aspects of physicochemical consistency, the rate at which the CMSI vent discharged boiling water (1–2 L second^-1^) fluctuated considerably. Dissolved inorganic carbon (DIC) concentration (3.5–5.6 mM), alongside the δ^13^C_DIC_ ratio (−2.53‰ to −1.9 ‰ V-PDB) which represented a low level of ^13^C depletion relative to the Vienna Pee Dee Belemnite standard (Table S1), also varied substantially. Furthermore, the solutes present in trace amounts (not >50 µM on any sampling occasion) exhibited high fluctuations in their concentrations, with iron, chromium, copper, and cobalt being the most inconsistent components of the vent-water. However, no diurnal or seasonal periodicity was associated with any of the fluxes mentioned above (Fig. 2). The DIC concentrations recorded for the vent-water were considerably high, compared to values reported from other hot springs. These numbers, together with the δ^13^C_DIC_ signatures, point towards a history of active circulation of the geothermal fluid through deep-seated limestone-/dolomite-like bedrocks, involving carbon dioxide dissolution via metamorphic degassing as well as microbial breakdown (respiration) of organic matter^28–30^.

### Temporal flux of cell density in the vent-water

Abundance of microbial cells in the vent-water was found to fluctuate widely over the six sampling occasions. Whereas total cell density ranged between 1.7×10^4^ mL^-1^ and 3.1×10^5^ mL^-1^, live cells remained between 1.1×10^4^ mL^-1^ and 8.2×10^4^ mL^-1^, and dead cell density ranged between 5.0×10^3^ mL^-1^ and 1.1×10^5^ mL^-1^ (Table 1). Every time the vent-water was sampled, proportion of live cells (55-75% of the total count recorded mL^-1^) exceeded that of dead cells (25-45% of the total count mL^-1^). Furthermore, on all the three exploration days, densities of both live and dead cells increased in the afternoon samples, compared with the corresponding forenoon samples; flux of live cells was always higher than that of dead cells, even as the magnitudes of the fluxes varied across the exploration days.

### Temporal consistency of the microbiome’s architecture

The six metagenomic analyses, undertaken over the six sampling occasions, collectively yielded 90 bacterial and 41 archaeal MAGs (Tables 1 and S2), which in turn clustered into 35 bacterial and 11 archaeal species (Table S3) on the basis of >95% orthologous average nucleotide identity, and >70% digital DNA-DNA hybridization, among the sibling MAGs (Table S4). Each of these species-level clusters was also discernible as robust branches within phylogenetic trees reconstructed based on universal bacterial (Fig. 3) or archaeal (Fig. 4) core genes. Among the sibling MAGs constituting a species, the largest one in terms of nucleotide length was chosen to represent the species in all downstream analyses (Table S2). For 26 out of the 46 microbial species identified, sibling MAGs got reconstructed on multiple sampling occasions (Table S3). Corroborative to this mark of microbiome consistency, 78-97% reads from each metagenomic dataset mapped individually onto the primary assemblies of the other five metagenomes (Table S5).

**Figure 3.**
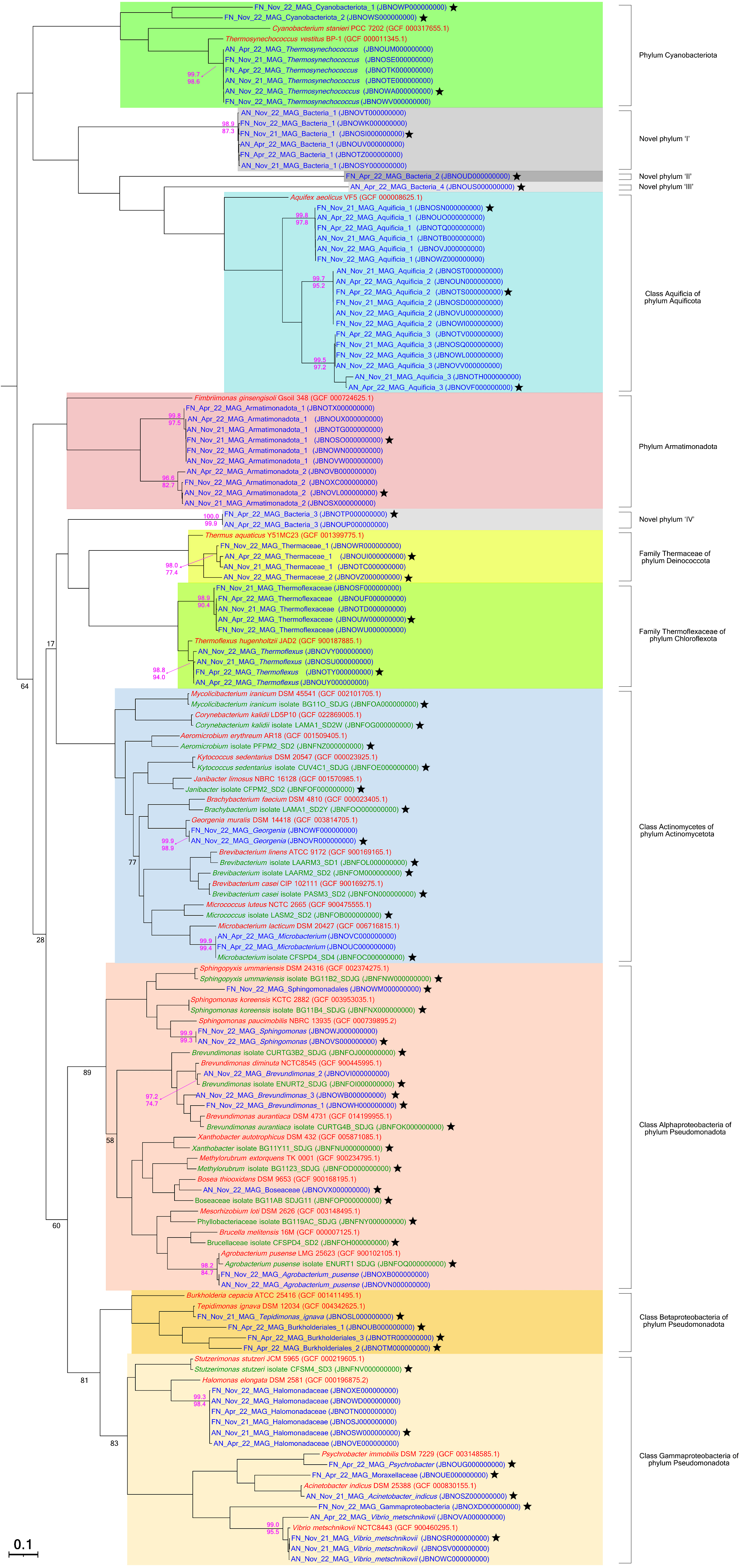
Maximum likelihood tree showing the evolutionary relationships existing among the 90 bacterial MAGs reconstructed over the six sampling occasions, the 23 bacterial species isolated on 01-Nov-2022, and the type strains of the taxa up to which the different MAGs or pure-culture isolates could be classified (in case a type strain’s genome sequence was unavailable, the genome of a strain nearest to the type was included in the analysis). The tree was constructed on the basis of the sequence similarities that the aforesaid entities possessed in relation to a conserved set of 92 universal bacterial core genes. The scale bar denotes a distance equivalent to 10% nucleotide substitution, deduced using the generalized time-reversible (GTR) model. Bootstrap values (with respect to 100 tests) recorded for all the nodes of the tree were >90, excepting those for which bootstrap values have been shown. The lowest pair-wise OrthoANI and GGDC values obtained among the entities clustered in a species-level branch have been indicated in pink font numerals at the root of the branch. Names of the MAGs, pure-culture isolates, and reference type strains are written in blue, green, and red fonts respectively. The major clades unifying the species-level clusters of MAG(s) and/or isolate(s) at the level of phylum (or class, within Pseudomonadota) are demarcated by different color shades. Every MAG/genome representing a species-level cluster in the downstream analyses is pointed out by a black star.

**Figure 4.**
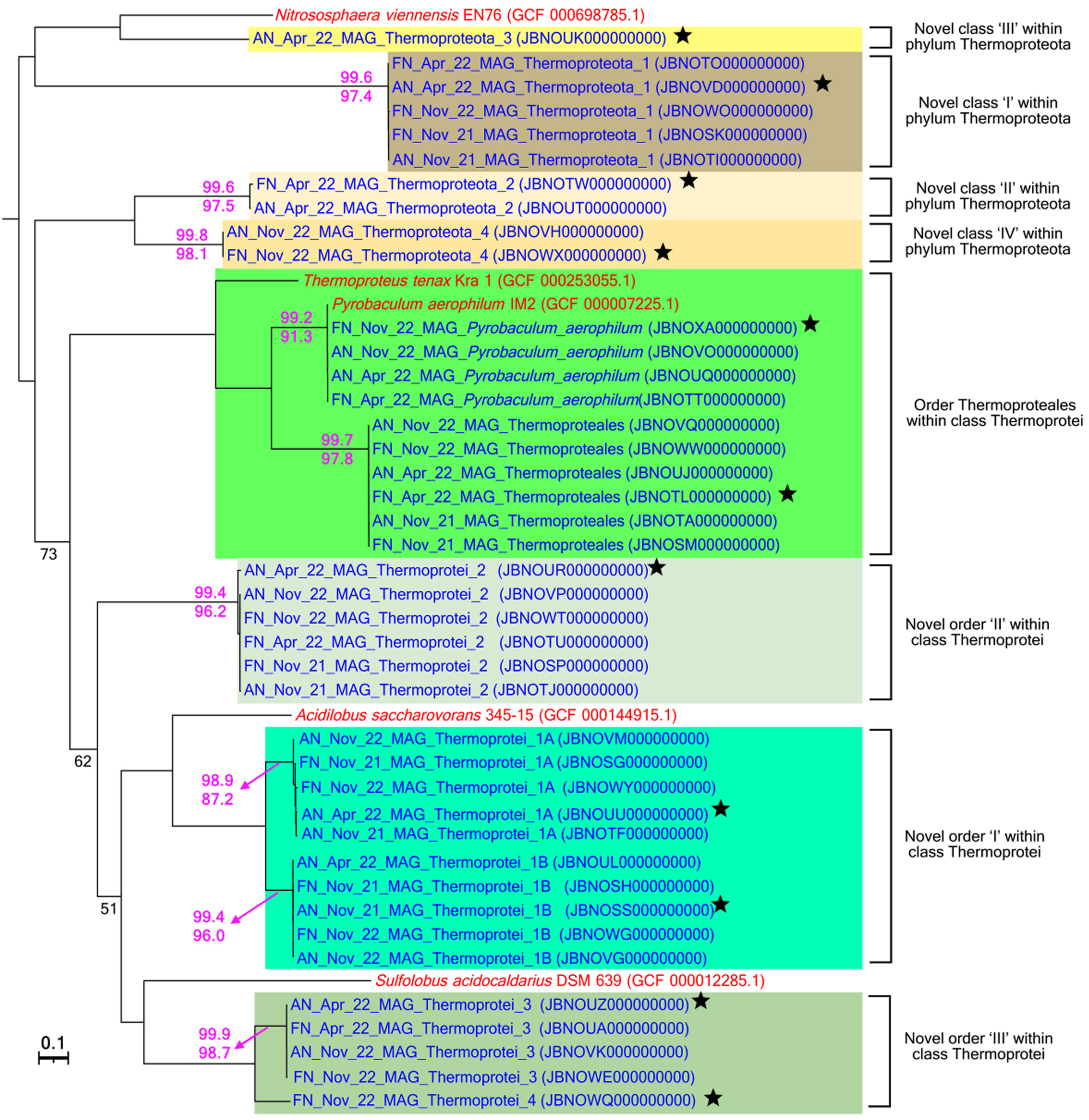
Maximum likelihood tree showing the evolutionary relationships existing among the 41 bacterial MAGs reconstructed over the six sampling occasions, and the type strains (or strains closely related to the types) of the taxa up to which the different MAGs could be classified. The tree was constructed on the basis of the sequence similarities that the aforesaid entities possessed in relation to a conserved set of 128 universal archaeal core genes. The scale bar denotes a distance equivalent to 10% nucleotide substitution, deduced using the generalized time-reversible (GTR) model. Bootstrap values (with respect to 100 tests) recorded for all the nodes of the tree were >90, excepting those for which bootstrap values have been shown. The lowest pair-wise OrthoANI and GGDC values obtained among the entities clustered in a species-level branch have been indicated in pink font numerals at the root of the branch. Names of the MAGs and reference type strains are written in blue and red fonts respectively. The major clades unifying the species-level clusters at the phylum level are demarcated by different color shades. MAGs representing the species-level clusters in downstream analyses are marked by black stars.

### Bacteria isolated from the boiling vent-water at 37°C

Of the 35 bacterial species identified as MAGs, six belonged to taxa that do not have any member strain known for laboratory growth at >45°C, whereas 14 belonged to such taxa which have only a few members reported for laboratory growth at >45°C, but not above 75°C (Table S3). Consequently, to determine whether these mesophiles and potential moderate thermophiles, or the likes of them, were alive *in situ*, enrichment and pure-culture isolation at 37°C, using 14 different culture media, were attempted upon the sample WD_Nov_22_Pc (Table 1; Fig. 1). Overall, 50 and 45 strains belonging to Actinomycetota and Pseudomonadota respectively got isolated on seven out of the 14 media types used (Table S6). Based on their 16S rRNA gene sequence similarities the actinobacterial and proteobacterial isolates could be clustered into 11 and 12 species respectively. Of the 11 Actinomycetota species isolated, five belonged to mesophilic taxa, and six were classified under taxa having few moderately-thermophilic members. Of the 12 Pseudomonadota species isolated, seven belonged to mesophilic taxa, whereas five were classified under taxa having few moderately-thermophilic members.

### Towards a holistic picture of population ecology involving all the uncultivated and cultivated CMSI species

Whole-genome shotgun sequence was determined for one representative strain from each species-level cluster of pure-culture isolates (Table S6). The assembled genomes of the 23 representative isolates obtained in this way were collated with the 46 representative MAGs already in hand to give rise to the **c**omprehensive **s**pecies-**l**evel **M**AG/**g**enome **s**equence (**CS**-**LMGS**) database for the CMSI vent-water microbiome (Table S7). Three species-level entities that were discovered as MAGs, and identified as *Agrobacterium_pusense*, *Brevundimonas*_2, and a member of *Microbacterium*, were also retrieved in the form of pure-cultures identified as *Agrobacterium pusense* isolate ENURT1_SDJG, *Brevundimonas* isolate ENURT2_SDJG, and *Microbacterium* isolate CFSPD4_SD4 (Fig. 3). Consequently, to avoid subject-sequence redundancy, for these three species, only the pure-culture genomes were included in the CS-LMGS database. To obtain a comprehensive picture of temporal population dynamics involving all the 66 species discovered, six different read-mapping experiments against the CS-LMGS database were performed using the six metagenomic datasets as individual queries (Fig. 5A). To estimate what proportion of reads from a query (metagenomic) dataset matched (aligned) spuriously with the MAGs/genomes included in the CS-LMGS database, the genome of a microbial species unlikely to be present in the CMSI vent-water (namely, the marine planktonic bacterium *Candidatus* Pelagibacter ubique HTCC1062, which belongs to the SAR11 clade of Alphaproteobacteria) was added to the database; this took the total number of species-level entities in the CS-LMGS database to 67 (Table S7).

**Figure 5.**
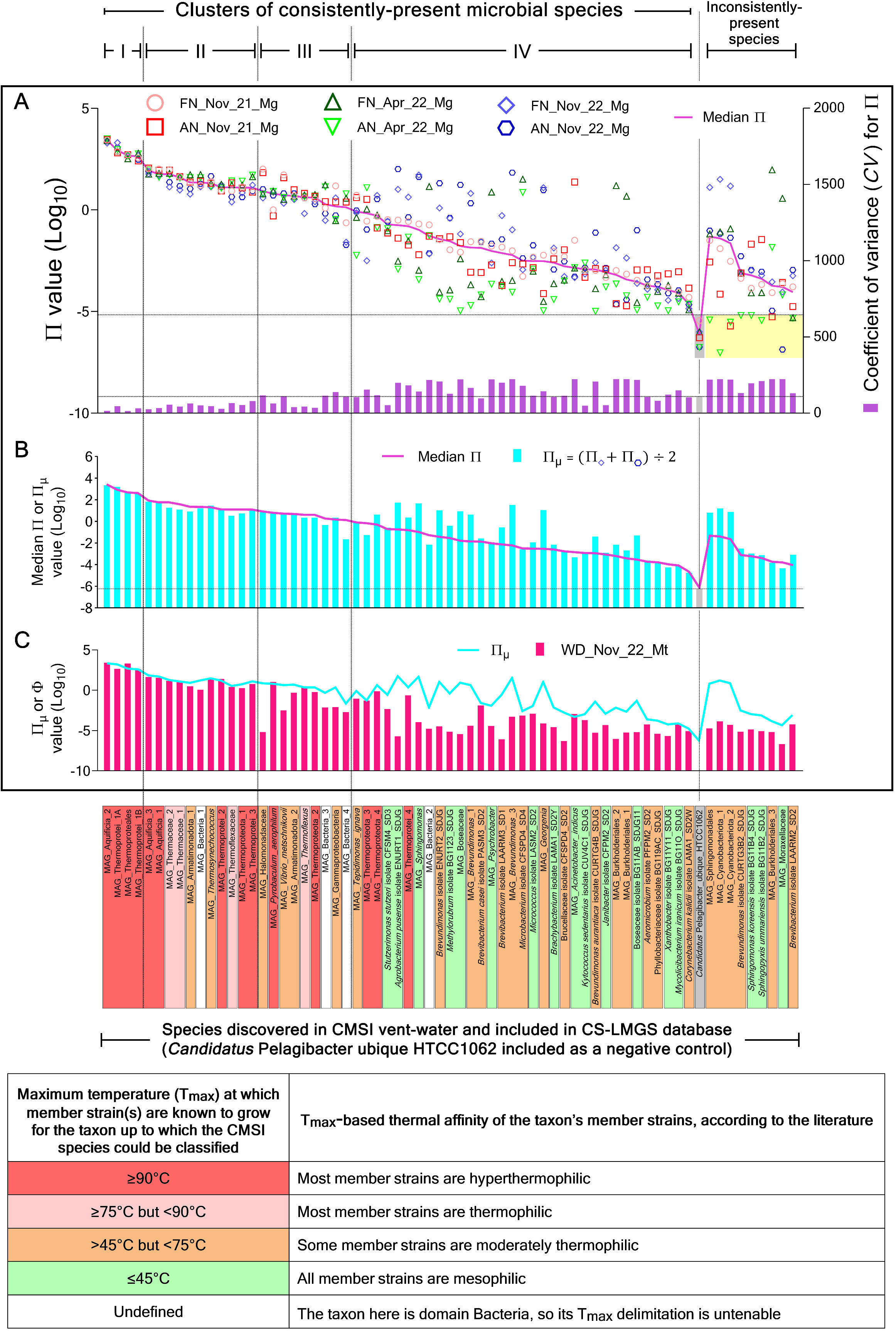
Coefficients of prevalence and functionality depicted for all the 66 microbial species that were discovered in the CMSI vent-water over the six sampling occasions; data obtained for the negative control *Candidatus* Pelagibacter ubique HTCC1062 have also been included. (**A**) For every species, all the individual Π values recorded over the six sampling occasions are plotted alongside the coefficient of variance (*CV*) determined for the fluctuation of Π. Π values below 1×10^-5^, which were considered to have resulted from spurious read-matching events, are demarcated by a yellow-shaded area. (**B**) For every species, its mean Π value recorded over the two sampling sessions of 01-Nov-2022 (Π_μ_) is plotted alongside the median of the six individual Π values (Π_median_) recorded over the six sampling occasions. (**C**) For each species, its Φ recorded through 01-Nov-2022 is plotted alongside Π_μ_. The consistently-present species are arranged according to the descending order, and clustered based on the ranges, of their Π_median_ values. The inconsistently-present species are clustered separately and arranged based on their own series of descending Π_median_ values. The data points plotted for *Candidatus* Pelagibacter act as the partition between the data obtained for the species present consistently and inconsistently. Uncultivated species are identified by the prefix “MAG” in their descriptions, whereas the cultured species have the term “isolate” within their descriptions. Species names are highlighted by different colors based on their putative thermal affinities (the name of the negative control is highlighted by a grey shade).

### Numeral supremacy of an Aquificia and a few Thermoprotei amid sustained presence of several other species

72-78% of the approximately 52-83 million high-quality reads contained in the different metagenomic datasets matched with sequences from the 66 CMSI MAGs/genomes included in the CS-LMGS database. Across the six “metagenome versus MAGs/genomes correspondence studies” conducted, negligible proportions (5×10^-5^ to 7×10^-6^ percent) of reads from the individual metagenomes matched sequences from *Candidatus* Pelagibacter ubique (Table S8). For the individual CMSI species, over the six metagenomic-read-mapping experiments, arithmetic mean of the average lengths recorded for aligned reads ranged between 148 and 238, whereas the mean recorded for the percentages of the MAG/genome length matched by aligned reads (coverage breadth) ranged between 0.4 and 100 (Fig. S1A).

With reference to a given sampling occasion, relative abundance of each species included in the CS-LMGS database was estimated as a coefficient of prevalence (Π), defined as that percentage of reads from the relevant metagenomic dataset which matched sequences from the MAG/genome of the species in question, per mega base length of the MAG/genome, multiplied by the MAG-/genome-wide coverage breadth achieved in the read mapping experiment. Across the six sampling occasions (metagenomic investigations), highest Π value recorded for *Candidatus* Pelagibacter ubique was 4×10^-6^. Consequently, 1×10^-5^ was considered to be the cut-off below which all coefficients of prevalence were regarded as results of spurious read-matching (see the data plotted within the yellow-shaded area of Fig. 5A). 57 CMSI species never had a Π value <1×10^-5^ so they were inferred to be present consistently in the habitat over the entire exploration period. Subsequently, based on their median Π values, the 57 **c**onsistently-**p**resent **m**icrobial **s**pecies (**CPMS**) could be classified into four distinct clusters. For the remaining nine species, Π was <1×10^-5^ on at least one sampling occasion, so they were not only inferred as virtually absent from the habitat on those instances, but also regarded, overall, as the inconsistently-present members of the microbiome (Fig. 5A).

1. The first **c**luster of **CPMS** (**CCPMS**-I) encompassed only four species, Aquificia_2, Thermoprotei_1A, a Thermoproteales member, and Thermoprotei_1B, characterized by >100 Π_median_ and overall low fluctuation of Π [coefficient of variance (*CV*) between 15 - 45].
2. CCPMS-II encompassed 11 members, each having Π_median_ between 10 and 100 (and *CV* of Π 26 - 80). Of these, two species each belonged to Aquificia, Thermaceae, and Thermoprotei, and one each belonged to Armatimonadota, Bacteria, Thermoflexaceae, Thermoproteota, and *Thermosynechococcus*.
3. CCPMS-III members had Π_median_ >1 but <10, alongside *CV* of Π 35 - 139. Out of the total nine species constituting this cluster, two could be classified only up to Bacteria, and one each belonged to Armatimonadota, Gammaproteobacteria, Halomonadaceae, *Pyrobaculum*, *Thermoflexus*, Thermoproteota, and *Vibrio*.
4. CCPMS-IV members had Π_median_ >2×10^-5^ but <1, in conjunction with *CV* of Π 50 - 223. Of the 33 species encompassing this cluster, 30 and three belonged to the domains Bacteria and Archaea respectively.
5. The **c**luster of **i**nconsistently-**p**resent **m**icrobial **s**pecies (**CIPMS**) encompassed nine members having Π_median_ between 2×10^-5^ and 5×10^-2^, but Π_minimum_ invariably <1×10^-5^ (*CV* of Π ranged from 131 to 224).

The patterns of species prevalence recorded over time in conjunction with the trends of vent-water chemistry identify CMSI as a stable geomicrobiological system. Whereas all the major biotic and abiotic components of the ecosystem remain steady over time, the rate of geothermal fluid discharge alongside a few minor components of the ecosystem, namely the trace solutes and sparse microbes present in the vent-water, are in states of uneven flux. This suggests that the system receives steady hydrological, and thereby microbiological, input from a major unwavering geological source, while a few minor sources might also make unstable contributions.

### *In situ* functionality of the vent-water species

74% of the 74 million high-quality mRNA-related reads contained in the bulked metatranscriptomic dataset WD_Nov_22_Mt (Table 1; Fig. 1) matched sequences from the 66 CMSI MAGs/genomes; not a single read aligned with any sequence from *Candidatus* Pelagibacter ubique. Furthermore, of the 55 million metatranscriptomic reads matching sequences from across the different CMSI MAGs/genomes, 96% were attributed to the four CCPMS-I members: 46% to Aquificia_2, 34% to the Thermoproteales species, and 8% each to Thermoprotei_1A and Thermoprotei_1B (Table S8). Fidelity of the metatranscriptome to the CMSI vent-water microbiome was corroborated by the fact that 95-96% reads from the metatranscriptomic dataset mapped onto the primary assemblies of the six different metagenomes sequenced (Table S5). For the individual CMSI species, average length recorded for aligned metatranscriptomic reads ranged between 20 and 99, whereas the percentage of protein-coding genes matched by metatranscriptomic reads (transcriptional breadth) ranged between 0.1 and 100 (Fig. S1B).

For a species included in the CS-LMGS database, coefficient of functionality (Φ), as on 01-Nov-2022, was determined as that percentage of reads from the metatranscriptomic dataset which matched sequences from its MAG/genome, per mega base length of the MAG/genome, multiplied by the MAG-/genome-wide transcriptional breadth achieved in the read mapping experiment. Φ varied not only across, but also within, the prevalence-based categories; however, Φ_maximum_ of a category decreased as follows: CCPMS-I > CCPMS-II > CCPMS-III > CCPMS-IV > CIPMS (Fig. 5C). For an objective interpretation of *in situ* functionality in the context of prevalence, Φ of a species was compared exclusively with the arithmetic mean of its forenoon and afternoon Π values (Π_μ_) recorded on 01-Nov-2022, i.e. the day the metatranscriptome was analyzed. Corroborative to this approach, Π_μ_ and Π_median_ values exhibited significant positive correlation across the 66 CMSI species (*R*^2^ = 0.90, *P* = 0.00), underscoring Π_μ_ as representative of the general trend of species prevalence existing in the habitat over time (Fig. 5B). Furthermore, Π_μ_ of none of the CMSI species was less than the 1×10^-5^ cut-off, below which all Π values were considered to be resulting from spurious read-matching (Table S8). Φ:Π_μ_ ratios (Table S8; Figs. 5C and S2) varied across, as well as within, the prevalence-based categories, although the two indices had significant positive correlation (*R*^2^ = 0.62, *P* = 0.00) across the 66 CMSI species.

### Differential expression of functional genes and metabolic systems across the species

Out of the 55 million metatranscriptomic reads matching sequences from across the 66 CMSI MAGs/genomes, 45 million mapped on to protein-coding genes (Tables S9 to S74). For every coding sequence (CDS) annotated in a given MAG/genome, its expression-level (EL_gene_) at the microbiome scale (Figs. 6 and 7) was delineated as the number of metatranscriptomic reads matching its sequence, per kilobase of its length, per million reads that aligned overall with CDSs from across the 66 CMSI species. Within a given MAG/genome, for every metabolic category defined by **c**lusters of **o**rthologous **g**enes (**COG**-category), a transcriptional breadth (^CC^tb) was first determined as that percentage of CDSs comprising the category which was matched by metatranscriptomic reads (Table S75). Subsequently, the expression-level of the COG-category (EL_cc_) was delineated as the product of its ^CC^tb and the median of all EL_gene_ values recorded under it in the form of rational numbers excepting zero.

**Figure 6.**
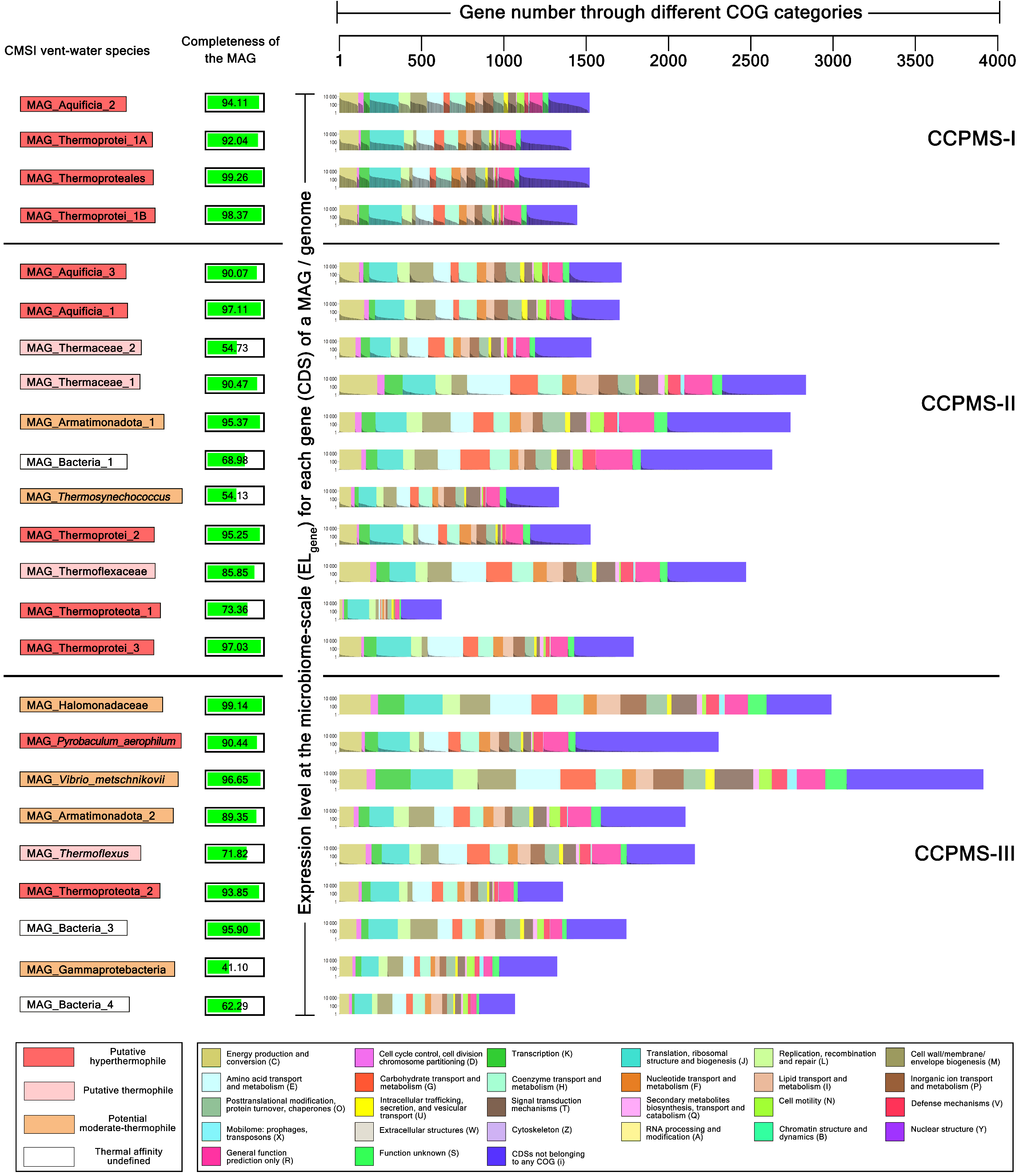
COG-category-wise distribution and microbiome-scale EL_gene_ values of the different CDSs annotated within the MAGs of the 24 CMSI species included in the first three clusters of consistently-present microbial species (CCPMS-I, II, and III). For the individual species, nomenclatures of the different CDSs annotated under the various COG-categories have been given Tables S9-S32 alongside the derivations of their EL_gene_ values. The 66 CMSI species have been arranged according their prevalence-based order defined in Fig. 5. Species names are highlighted by different colors based on their putative thermal affinities (the color code is same as the one used in Fig. 5).

**Figure 7.**
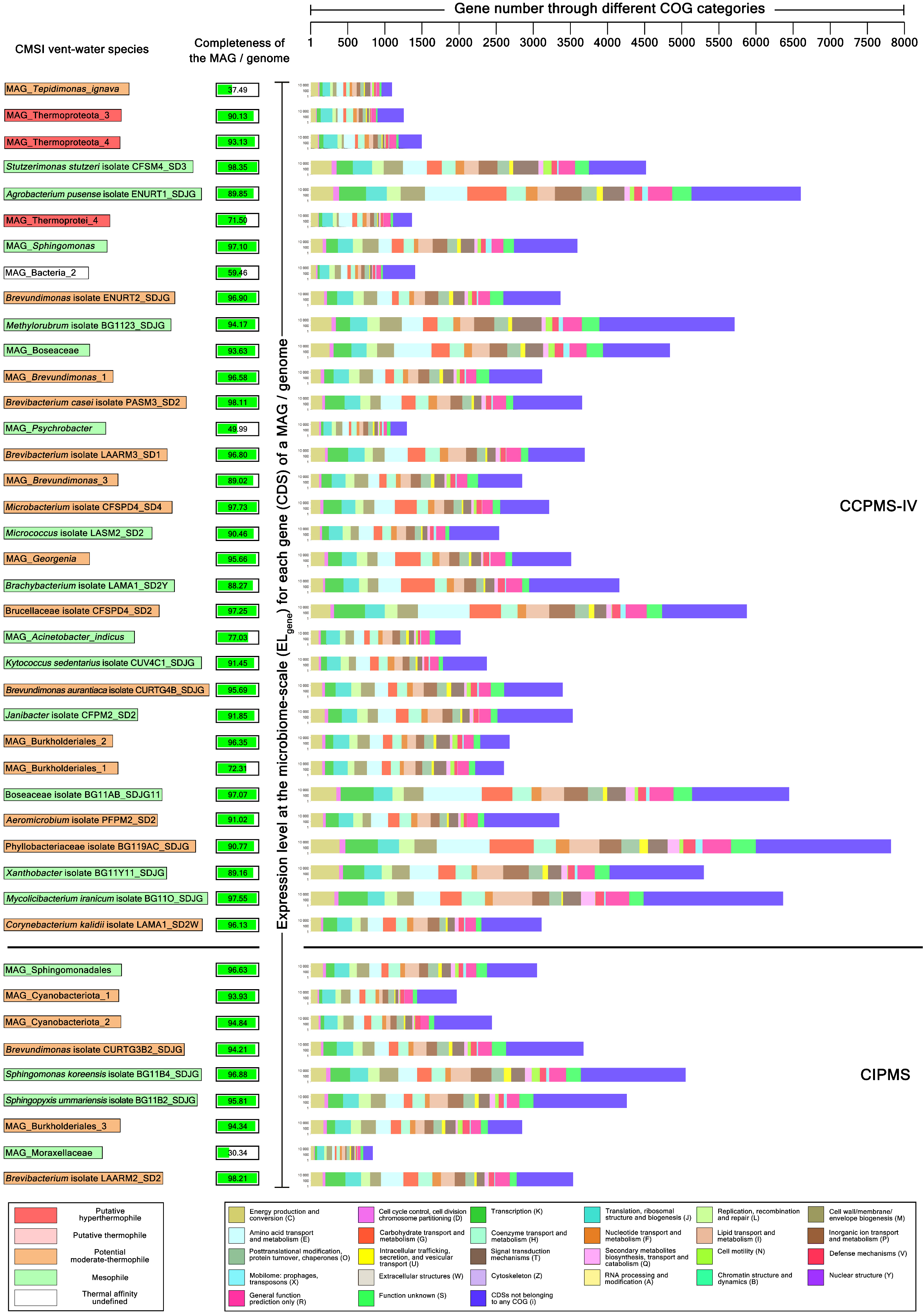
COG-category-wise distribution and microbiome-scale EL_gene_ values of the different CDSs annotated within the MAGs of the 42 CMSI species included in the fourth cluster of consistently-present microbial species (CCPMS-IV) and the cluster of inconsistently-present microbial species (CIPMS). For the individual species, nomenclatures of the different CDSs annotated under the various COG-categories have been given Tables S33-S74 alongside the derivations of their EL_gene_ values. The 66 CMSI species have been arranged according their prevalence-based order defined in Fig. 5. Species names are highlighted by different colors based on their putative thermal affinities (the color code is same as the one used in Figs. 5 and 6).

17 CMSI species had up to 100% ^CC^tb for every COG-category that was encompassed by their MAGs in relation to cell system maintenance and growth (COG-categories represented by the symbols C, D, K, J, L, and M), structural aspects of cell system maintenance (E, G, H, F, I, and P), and environmental responses of the cells (O, U T, Q, N, and V; COG-category abbreviation code given in Figs. 6 and 7). These species included all the four members of CCPMS-I; all CCPMS-II members except Armatimonadota_1; and the CCPMS-III members Armatimonadota_2, *Pyrobaculum*_*aerophilum*, and *Thermoflexus* species (Fig. 6). Within this grouping, and compared to any other CMSI species as well, CCPMS-I members had much higher EL_cc_ for all the 18 physiologically-fundamental COG-categories (PFCCs) mentioned above (Fig. 8). However, at the levels of the biochemical pathways, structural complexes, and functional units defined in BRITE Tables (Fig. 9) and KEGG Modules (Table S76; Fig. 10), all 17 species of this grouping were unified by the expression of genes central to cell division (Fig. 9), besides showing metatranscriptomic signatures for aminoacyl-tRNA synthetases, DNA and RNA polymerases, ribosomal proteins (Fig. 9), and 11 basic metabolic modules. Excluding Thermoproteota_1, for which MAG-length was only 563 kilobases, all members of this group exhibited metatranscriptomic signatures for a common set of 24 more KEGG Modules (Fig. 10).

**Figure 8.**
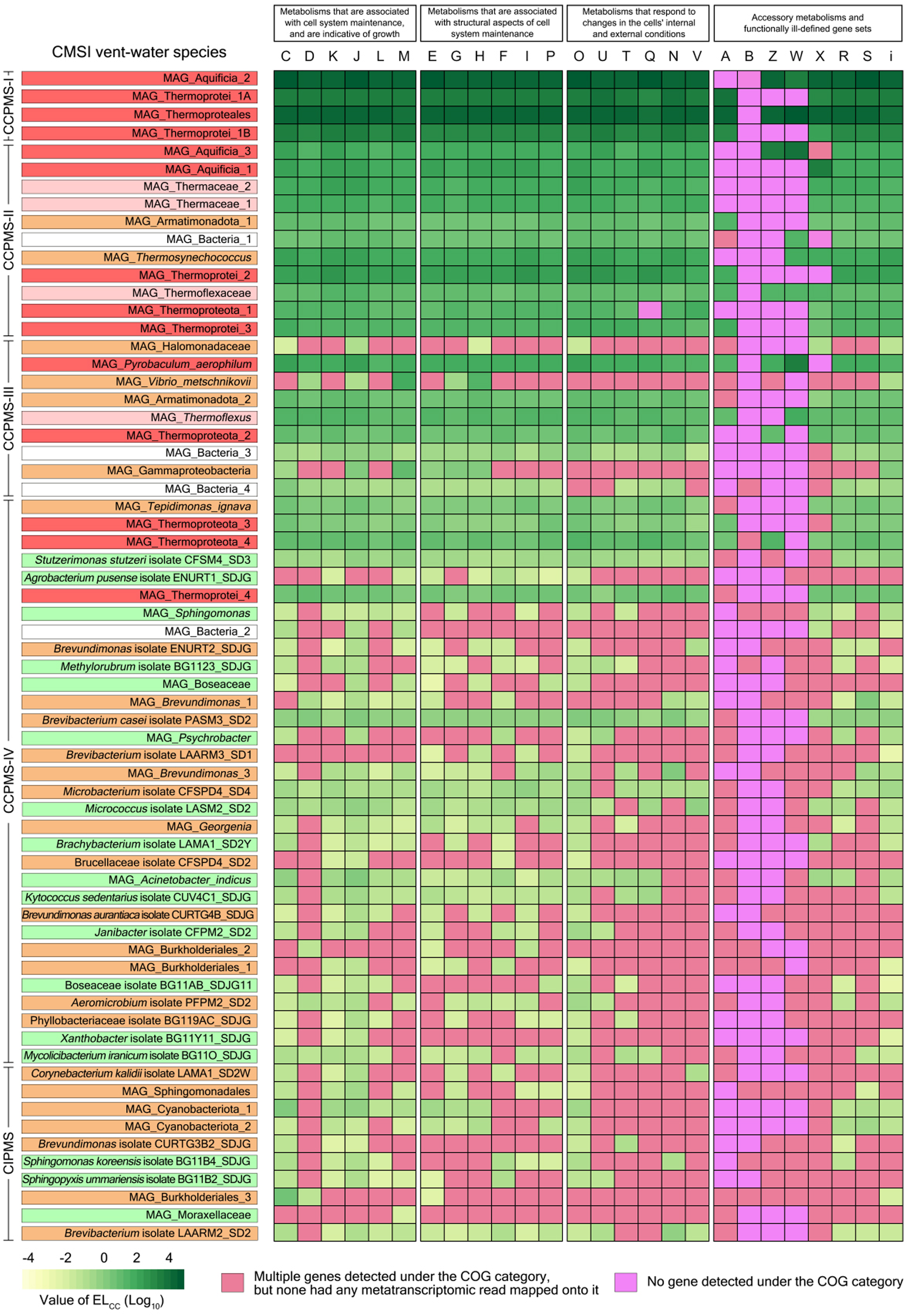
Heat map comparing the *in situ* expression-levels of the different COG-categories (EL_cc_) that were detected within the MAGs/genomes of the 66 CMSI species. The yellow to green cell-color gradient shown in the bottom left corner of the figure tracks the increase in EL_cc_ across COG-categories as well as species. Absence of a COG-category within a MAG/genome is depicted by a pink cell, whereas the absence of expression signature for a COG-category present within a MAG/genome is depicted by a red cell. For the individual species, Abbreviation code used for the different COG-categories is same as that shown in Figs. 6 as well as 7. Derivation and numeral magnitude of each EL_CC_ value underlying the heat-map are given in Table S75. The 66 CMSI species have been arranged according their prevalence-based order defined in Fig. 5. Species names are highlighted by different colors based on their putative thermal affinities (the color code is same as the one used in Figs. 5 through 7).

**Figure 9.**
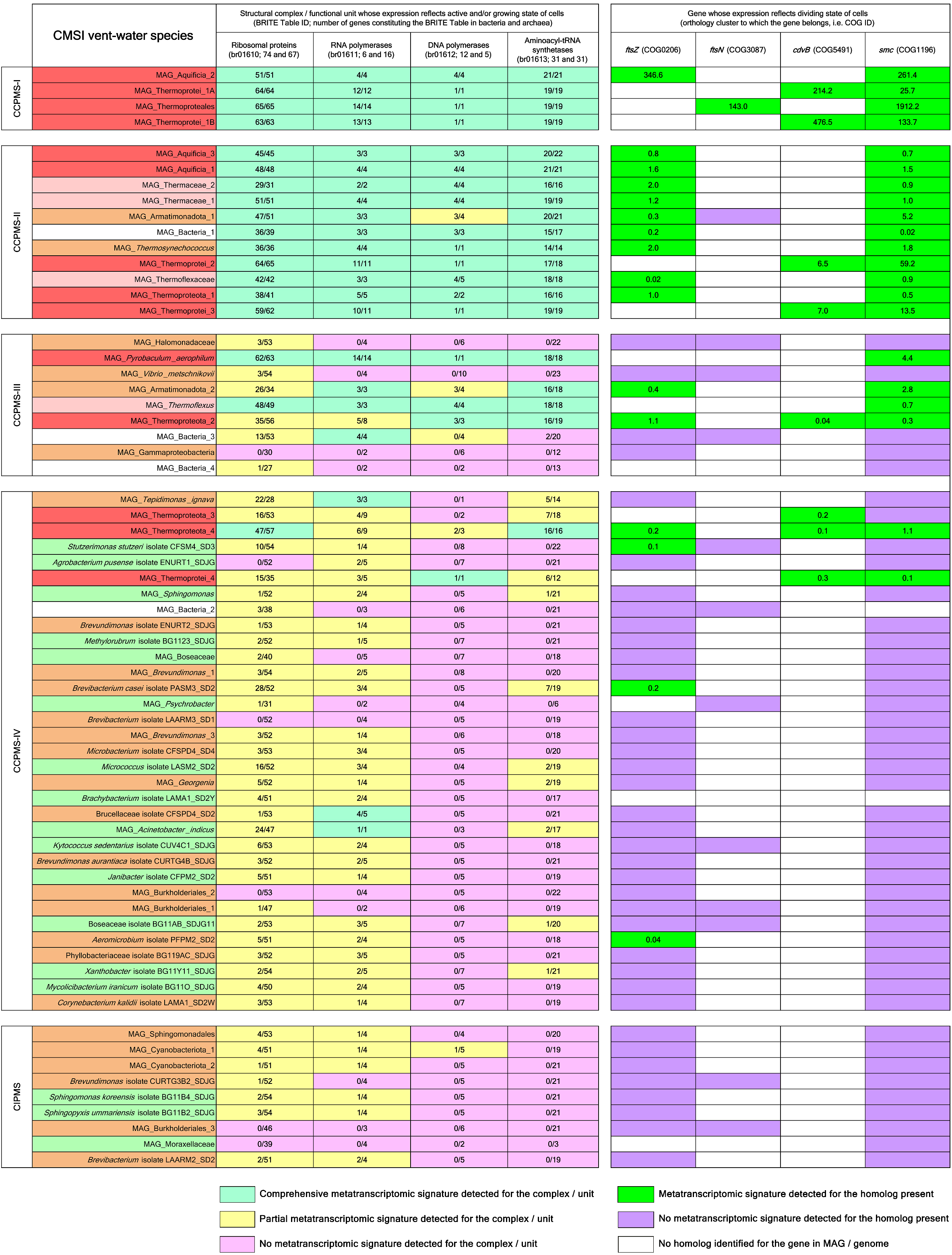
*In situ* expression of structural complexes and functional units (BRITE Tables), or individual genes, the transcription of which potentially reflect active and/or growing state of cells. Comprehensive metatranscriptomic signature was said to have been detected for a BRITE Table when ≥80% of the genes detected for the Table in the MAG/genome were expressed. Partial metatranscriptomic signature was said to have been detected for a BRITE Table when <80% of the genes detected for the Table in the MAG/genome were expressed. No metatranscriptomic signature was said to have been detected for a BRITE Table when none of the genes detected for the Table in the MAG/genome were expressed. The 66 CMSI species have been arranged according their prevalence-based order defined in Fig. 5. Species names are highlighted by different colors based on their putative thermal affinities (the color code is same as the one used in Figs. 5 through 8).

**Figure 10.**
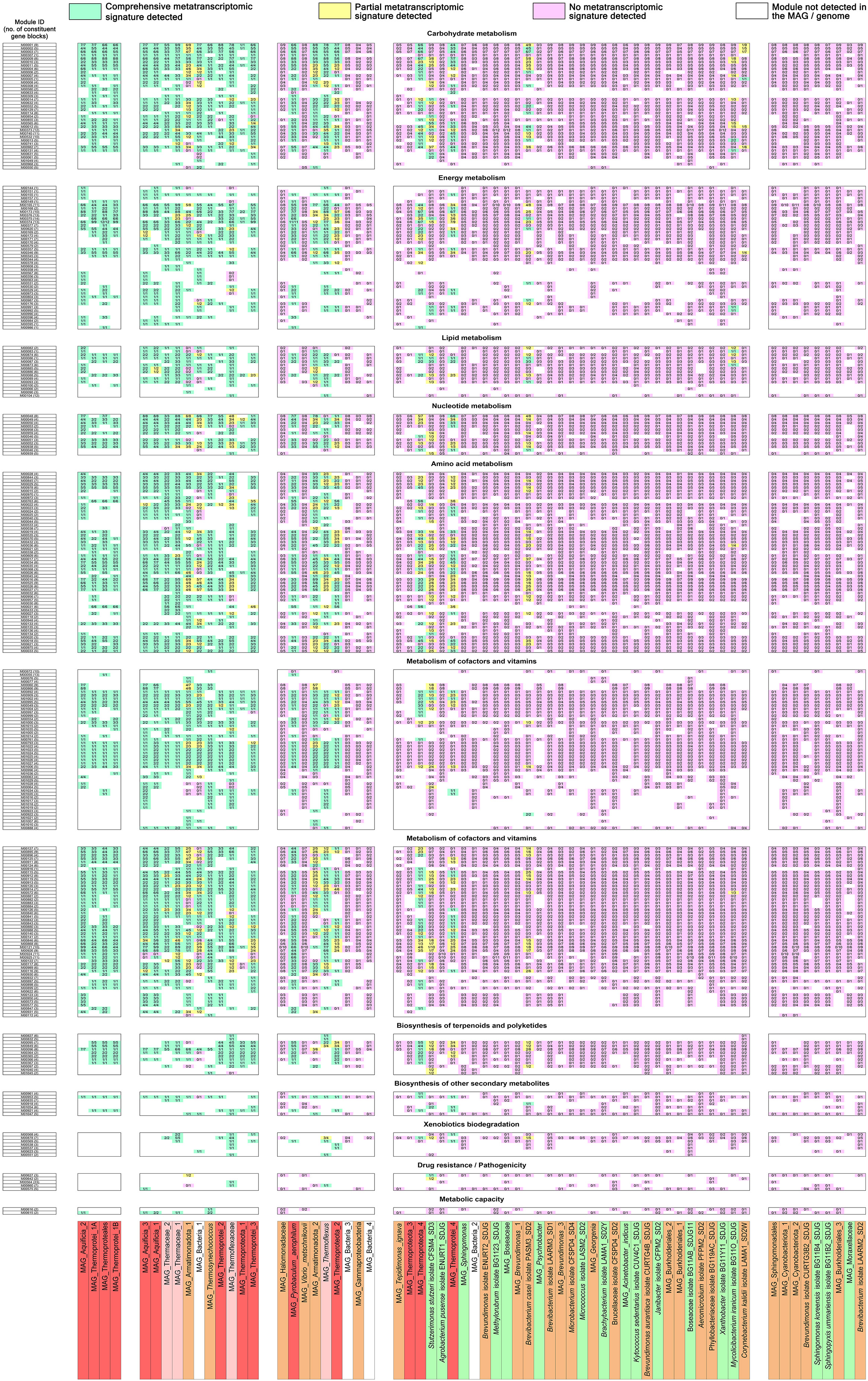
Biochemical pathways, structural complexes, and functional units (KEGG Modules) expressed by the different CMSI vent-water species *in situ*. Comprehensive metatranscriptomic signature was said to have been detected for a KEGG Module when ≥80% of its gene-blocks detected completely within the concerned MAG/genome were expressed in their entirety. Partial metatranscriptomic signature was said to have been detected for a KEGG Module when <80% of its gene-blocks detected completely within the concerned MAG/genome were expressed in their entirety. No metatranscriptomic signature was said to have been detected for a Module when none of the gene-blocks detected completely for the module in the MAG/genome were expressed in their entirety. A Module was said to be missing from a MAG/genome when none of its known gene-blocks were detected completely. All the 289 Modules expressed by the different CMSI species have been indicated here by their KEGG IDs; complete biochemical descriptions can be found in Table S76. The 66 CMSI species have been arranged according their prevalence-based order defined in Fig. 5. Species names are highlighted by different colors based on their putative thermal affinities (the color code is same as the one used in Figs. 5 through 9).

Nine CMSI species had up to 90% ^CC^tb for each of the 18 PFCCs; they included Armatimonadota_1 from CCPMS-II; Bacteria_3 and Thermoproteota_2 from CCPMS-III (Fig. 6); and the CCPMS-IV members *Brevibacterium casei* isolate PASM3_SD2, *Stutzerimonas stutzeri* isolate CFSM4_SD3, *Tepidimonas*_*ignava*, Thermoprotei_4, Thermoproteota_3, and Thermoproteota_4 (Fig. 7). Among these species, Armatimonadota_1, Thermoprotei_4, Thermoproteota_2, and Thermoproteota_4 had higher EL_cc_ for all the 18 PFCCs, compared to the remaining five entities (Fig. 8); they also expressed a number of genes central to cell division whereas the other five did not (Fig. 9). All the nine species of this grouping showed metatranscriptomic signatures for ribosomal proteins and RNA polymerases (Fig. 9). Armatimonadota_1, Thermoprotei_4, Thermoproteota_2, Thermoproteota_4, *Brevibacterium casei* PASM3_SD2, and *Stutzerimonas* CFSM4_SD3, exhibited signatures for a common set of 26 KEGG Modules (Fig. 10), whereas Bacteria_3, *Tepidimonas*_*ignava*, and Thermoproteota_3 showed none.

Bacteria_4 from CCPMS-III (Fig. 6); *Acinetobacter*_*indicus*, *Brevundimonas*_3, the *Georgenia* species, *Kytococcus sedentarius* isolate CUV4C1_SDJG, *Microbacterium* CFSPD4_SD4, *Micrococcus* isolate LASM2_SD2, and *Mycolicibacterium iranicum* isolate BG11O_SDJG from CCPMS-IV; and *Brevibacterium* isolate LAARM2_SD2 from CICPMS (Fig. 7), exhibited up to 20% ^CC^tb across 12 or more PFCCs each. Overall, these nine species had low EL_cc_ across COG-categories, but expression-levels for Energy production and conversion; Translation, ribosomal structure and biogenesis; and Inorganic ion transport and metabolism were relatively higher (Fig. 8). All the nine species showed metatranscriptomic signatures for ribosomal proteins (Fig. 9). However, only *Mycolicibacterium* BG11O_SDJG was found to exhibit metatranscriptomic signatures for KEGG Modules (this species showed signatures for 17 modules in all; Fig. 10). No member of this grouping showed any metatranscriptomic signature for any key cell division gene (Figure 9).

The remaining 31 CMSI species (Figs. 6 and 7) had up to 15% ^CC^tb, and unequivocally low EL_cc_, across the few PFCCs that they expressed (Fig. 8). However, compared to the other COG-categories, EL_cc_ values were relatively higher for Energy production and conversion; Transcription; Translation, ribosomal structure and biogenesis; and Posttranslational modification, protein turnover, chaperones. 25 of these species showed metatranscriptomic signatures for ribosomal proteins, whereas 19 showed metatranscriptomic signatures for RNA polymerases (Fig. 9); only *Corynebacterium kalidii* isolate LAMA1_SD2W was found to exhibit metatranscriptomic signatures for KEGG Modules (this species showed signatures for seven modules in all; Fig. 10); in addition, only *Aeromicrobium* isolate PFPM2_SD2 exhibited metatranscriptomic signature for a key cell division gene, namely *ftsZ* (Fig. 9).

### The geothermal vent-water as an inclusive ecosystem, albeit with wide inter-community asymmetries

Out of the total 66 microbial species discovered, four could not be classified below the level of the domain Bacteria, whereas 62 were ascribable to 36 bacterial (Fig. 3) and four archaeal (Fig. 4) taxa ranging from the level of genus to that of phylum. These 62 species, according to *in vitro* growth literature, could be putatively ordered under four temperature-affinity-based communities (Tables S3 and S6; Fig. 5).

- 14 species were affiliated to five such taxa (Aquificia, *Pyrobaculum*, Thermoprotei, Thermoproteales, and Thermoproteota) that encompass mostly hyperthermophilic members. Of the total quanta of prevalence (cumulative Π) recorded for the 66 CMSI species, across the six different sampling occasions, 91–96% were attributed to these putative hyperthermophiles. Likewise, 99% of the total functionality (cumulative Φ) recorded for all CMSI species through 01-Nov-2022 was ascribed to these 14 species.
- Four species were affiliated to three such taxa (Thermaceae, Thermoflexaceae, and *Thermoflexus*) that encompass mostly thermophilic members. Of the cumulative Π and Φ values recorded, 0.5–3.3% and 0.52% were attributed to these putative thermophiles, respectively.

The 11 archaeal and three bacterial species ascribed to taxa of hyperthermophiles, together with the four bacterial species affiliated to taxa of thermophiles, were present in the CMSI vent-water consistently over time (Fig. 5A). Across these 18 species, *in situ* prevalence and functionality (Figs. 5C and S2) had a very strong positive relationship, with Φ and Π_μ_ values showing significant positive correlation across the 14 putatively hyperthermophilic (*R*^2^= 0.56, *P* = 0.002), as well as the four putatively thermophilic (*R*^2^ = 0.98, *P* = 0.006), species.

- 27 species were affiliated to 17 such taxa which encompass a few moderately-thermophilic members each. Of the cumulative Π and Φ values recorded, 1–5% and 0.46% were ascribed to these potential moderate-thermophiles, respectively.
- 17 species were affiliated to 15 such taxa that encompass only mesophilic members. Of the cumulative Π and Φ values recorded, 0.002–3% and 0.02% were attributed to these mesophiles respectively.

Of the 27 and 17 potential moderate-thermophiles and mesophiles discovered, 21 and 14 were always present in the habitat, respectively (Fig. 5A). Although occasionally absent from the vent-water, a few entities such as Cyanobacteriota_1, Cyanobacteriota_2, Burkholderiales_3, and a Sphingomonadales species, were at times prevalent at levels surpassing, or equivalent to, those of a number of putative hyperthermophiles (e.g. Thermoprotei_4 and Thermoproteota_3) and thermophiles (e.g. the *Thermoflexus* species and Thermaceae_1). Φ and Π_μ_ values (Figs. 5C and S2) showed weak but significant positive correlation across the potential moderate-thermophiles (*R*^2^ = 0.31, *P* = 0.003), and very weak and insignificant negative correlation across the mesophiles (*R*^2^ = 0.02, *P* = 0.6).

In the context of inter-community asymmetry, it is interesting to observe that the *Thermosynechococcus* species had higher Φ than seven, and higher Π_μ_ than eight, CMSI species classified under hyperthermophilic taxa (Fig. 5C). The *Thermosynechococcus* also had higher Φ as well as Π_μ_ than all the four putative thermophiles discovered. Likewise, the *Thermoflexus* and Thermoflexaceae species, together with Armatimonadota_1, had higher Φ and Π_μ_ than quite a few putative hyperthermophiles (Fig. 5C).

### Asymmetries within the temperature-affinity-based communities

Albeit 14 putative hyperthermophiles were detected in all, Aquificia_2, Thermoprotei_1A, the Thermoproteales species, and Thermoprotei_1B formed a core caucus of ecosystem dominance (Figs. 5 and S2). Furthermore, within this caucus of dominance, Aquificia_2 maintained absolute numeral supremacy throughout the exploration period, except in the forenoon of 01-Nov-2022 when its Π was marginally overtaken by that of Thermoprotei_1A (Table S8; Fig. 5A). Of the six individual cumulative Π values recorded, 86–93% were attributed to these four species, with Aquificia_2 alone accounting for 37–59%. Likewise, of the cumulative Φ, 97% was ascribed to the four predominant hyperthermophiles, with Aquificia_2 alone accounting for 45% (Table S8; Fig. 5C).

The 10 putative hyperthermophiles, which were not part of the preponderance caucus, accounted for only 3–5% and 2% of the cumulative Π and Φ values recorded. Furthermore, among these 10 species, only Aquificia_1 and Aquificia_3 jointly accounted for 2–4% and 1% of the cumulative Π and Φ values. The molecular attributes that may be instrumental to the overwhelming preponderance of Aquificia_2, Thermoprotei_1A, the Thermoproteales species, and Thermoprotei_1B have been delineated in Supplementary Results and Discussion, along with the plausible shortcomings that might cause the numeral and functional laggardness of the ecologically less-efficient archaea.

Of the six individual cumulative Π values recorded, 0.4-2.7% were attributed to Thermaceae_1 and Thermaceae_2, whereas only 0.1-0.6% were attributed to the Thermoflexaceae and *Thermoflexus* species; likewise, 0.4-1.5% were attributed to the *Thermosynechococcus* species and Armatimonadota_1, whereas all the other moderate-thermophiles and mesophiles collectively accounted for 0.4-5.9%. Of the cumulative Φ, 0.43% was attributed to the two Thermaceae species, whereas only 0.09% was attributed to the Thermoflexaceae and *Thermoflexus* species; likewise, 0.45% was attributed to the *Thermosynechococcus* species and Armatimonadota_1, whereas all the other moderate-thermophiles and mesophiles collectively accounted for 0.03%. The plausible molecular strategies underlying the ecological fitness of the putative thermophiles and potential moderate-thermophiles have been delineated in Supplementary Results and Discussion, along with the probable molecular drivers of the persistence exhibited by the CMSI mesophiles.

### Metabolic status of species across the temperature-affinity-based communities

All the putative hyperthermophiles except Thermoproteota_3; all the four putative thermophiles; the potential moderate-thermophiles Armatimonadota_1, Armatimonadota_2, and the *Thermosynechococcus* species; plus Bacteria_1, exhibited metatranscriptomic signatures for DNA and RNA polymerases, ribosomal proteins, aminoacyl-tRNA synthetases (Fig. 9), the chromosome segregation ATPase Smc^31^, plus at least one of the following markers of cell-division: (i) the GTPase FtsZ, (ii) FtsN which activates septal peptidoglycan synthesis and cell constriction, and (iii) CdvB that renders membrane constriction^32–34^. Hence, these 21 species are inferred to be growing (Fig. 11) *in situ*. 16 potential moderate-thermophiles, 13 mesophiles, Thermoproteota_3, and Bacteria_4, exhibited metatranscriptomic signatures for RNA polymerases and ribosomal proteins. However, out of these 31 species, only 11 showed metatranscriptomic signatures for DNA polymerases or aminoacyl-tRNA synthetases, four exhibited the same for FtsZ or CdvB, but none showed expression signatures for Smc (Fig. 9). Consequently, it seems that populations of these species do not grow (Fig. 11), despite considerable metabolic activities (Figs. 8 and 10), *in situ*.

**Figure 11.**
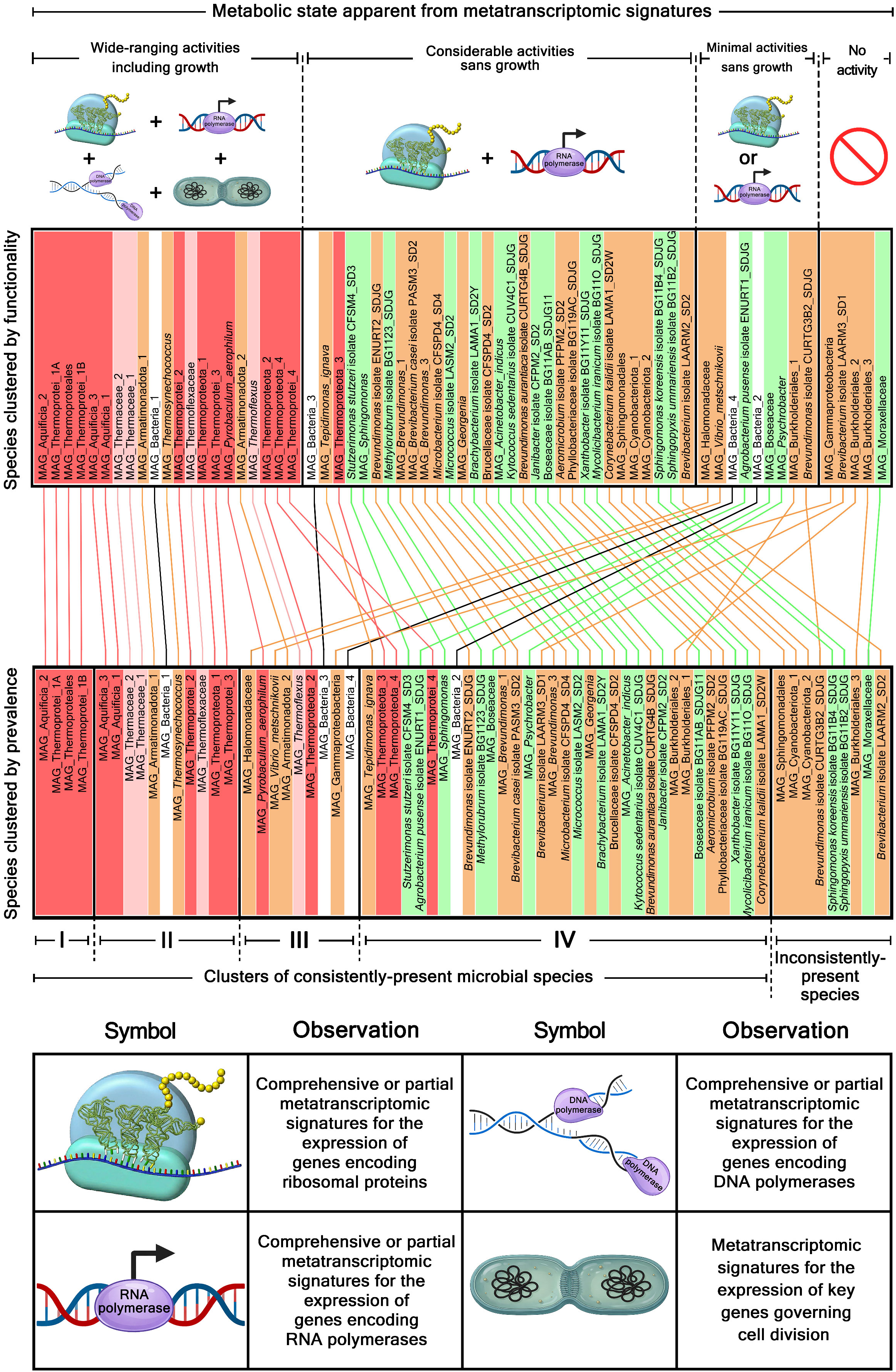
Graphical representation of which CMSI species are putatively (i) growing, (ii) metabolically active but not growing, (iii) barely alive (rendering extremely-limited activity), or (iv) metabolically inactive (dead for all practical purposes), *in situ*. This information pertaining to the eventual metabolic status of the species has been presented in the context of the prevalence-based status of the species. All uncultivated species have been identified by the prefix “MAG” in their descriptions, whereas the cultured species contain the term “isolate” within their descriptions. Species names are highlighted by different colors based on their putative thermal affinities (the color code is same as the one used in Figs. 5 through 10).

Four potential moderate-thermophiles, three mesophiles, Bacteria_2, and Bacteria_3, exhibited metatranscriptomic signatures for RNA polymerases or ribosomal proteins, but not DNA polymerases, aminoacyl-tRNA synthetases, Smc, FtsZ, FtsN, or CdvB (Fig. 9). Hence, populations of these nine species appear to render minimal metabolic activities, sans growth (Figs. 8, 10, and 11).

Four potential moderate-thermophiles, and one mesophile, showed no metatranscriptomic signature for DNA or RNA polymerases, ribosomal proteins, aminoacyl-tRNA synthetases, Smc, FtsZ, FtsN, or CdvB (Fig. 9). Therefore, from the metatranscriptomic point of view, populations of these five bacteria seemed to be inactive *in situ* (Figs. 8, 10, and 11). That said, isolation of one of these five species as a pure-culture (*Brevibacterium* strain LAARM3_SD1, which had a Π_μ_ of 0.3 and a Π_median_ of 0.01) pointed out that the absence of key metatranscriptomic signatures, or for that matter a Φ as low as 10^-6^, do not necessarily mean that the population is fully devoid of live cells *in situ*.

### (Hyper)thermophiles as primary producers of the vent-water ecosystem

Chemolithotrophy based on the oxidation of carbon monoxide, hydrogen, and reduced sulfur species was determined as the key energy-yielding metabolism of the CMSI microbiome, rendered exclusively by the putative hyperthermophiles and thermophiles. Metatranscriptomic reads (Table S77) corresponding to genes encoding the

- molybdenum-containing carbon monoxide dehydrogenase CoxL were detected for Aquificia_1, *Pyrobaculum*_*aerophilum*, Thermoprotei_2, Thermoprotei_3, Thermoprotei_4, Thermoproteota_2, Thermoproteota_4, and the two species classified under Thermoflexaceae and *Thermoflexus*;
- different subunits of [NiFe]-dependent hydrogenase, namely HyaA, HyaB, HyaC and/or HyaD, were detected for Aquificia_1, Thermoprotei_1B, Thermoprotei_2, and the two species affiliated to Thermoflexaceae and *Thermoflexus*;
- sulfide:quinone oxidoreductase SQOR were detected for Aquificia_1, Aquificia_2, Aquificia_3, *Pyrobaculum*_*aerophilum*, Thermaceae_1, Thermaceae_2, Thermoproteales, Thermoprotei_1B, Thermoprotei_3, Thermoprotei_4, Thermoproteota_4, and the *Thermoflexus* species;
- core subunits of the sulfur oxidation (Sox) system, namely SoxX, SoxA, SoxY, SoxZ, and SoxB, were detected for Aquificia_1, Aquificia_2, Aquificia_3, Thermaceae_1, and Thermaceae_2;
- intracellular sulfur oxidation protein DsrE/DsrF were detected for Aquificia_1, Aquificia_2, and Aquificia_3;
- thiosulfate dehydrogenase TsdA were detected for Aquificia_2, Thermaceae_1, Thermaceae_2, and the *Thermoflexus* species.

Reverse tricarboxylic acid (rTCA) cycle, modified rTCA cycle, and dicarboxylate/4-hydroxybutyrate cycle, were the main autotrophic carbon-fixation pathways employed by the putative (hyper)thermophiles for biomass generation (Table S77).

- Metatranscriptomic reads corresponding to some or all of the genes encoding (i) ATP-citrate lyase AclAB, (ii) NADH-dependent fumarate reductase FrdABCDE, and (iii) ferredoxin-dependent 2-oxoglutarate synthase/oxidoreductase KorABCD, which catalyze the key rTCA cycle reactions (i) citrate to acetyl-CoA and oxaloacetate, (ii) fumarate to succinate, and (iii) CoA to 2-oxoglutarate, respectively were detected for all the 14 putatively hyperthermophilic bacteria and archaea present in the vent-water.
- Metatranscriptomic reads corresponding to all the genes encoding (i) citryl-CoA synthetase and (ii) citryl-CoA lyase, which catalyze the two key reactions of the modified rTCA cycle, namely (i) citrate to citryl-CoA, and (ii) citryl-CoA to oxaloacetate and acetyl-CoA, respectively were detected for all the three putatively hyperthermophilic bacteria present in the vent-water.
- Metatranscriptomic reads corresponding to the gene encoding 4-hydroxybutyryl-CoA dehydratase, which governs the key step of converting 4-hydroxybutyryl-CoA to crotonyl-CoA in the dicarboxylate/4-hydroxybutyrate cycle, were detected for *Pyrobaculum*_*aerophilum*, Thermoproteales, and Thermoproteota_3.

### Ecological roles of the vent-water mesophiles and moderate-thermophiles

Mapping of metatranscriptomic reads on to the genomes of the CMSI mesophiles and potential moderate-thermophiles revealed their plausible ecological roles (Table S78; Fig. 12A) as scavengers of reactive oxygen species (ROS) and heavy metals, which can jeopardize overall microbiome health if not removed from the environment^35,36^. For instance, metatranscriptomic reads corresponding to some or all of the

i. peroxides-/superoxides-detoxifying genes *ahpC, ahpF*, *osmC*, *katE*, *katG* and *sodA* were detected for three mesophiles and eight potential moderate-thermophiles; whereas the
ii. heavy-metals-detoxifying genes *acr3*, *arsA*, *arsB*, *arsR*, *cadD*, *copC*, *copZ*, *czcD*, *fur*, *zntA*, and *znuA* were detected for four mesophiles and six potential moderate-thermophiles.

**Figure 12.**
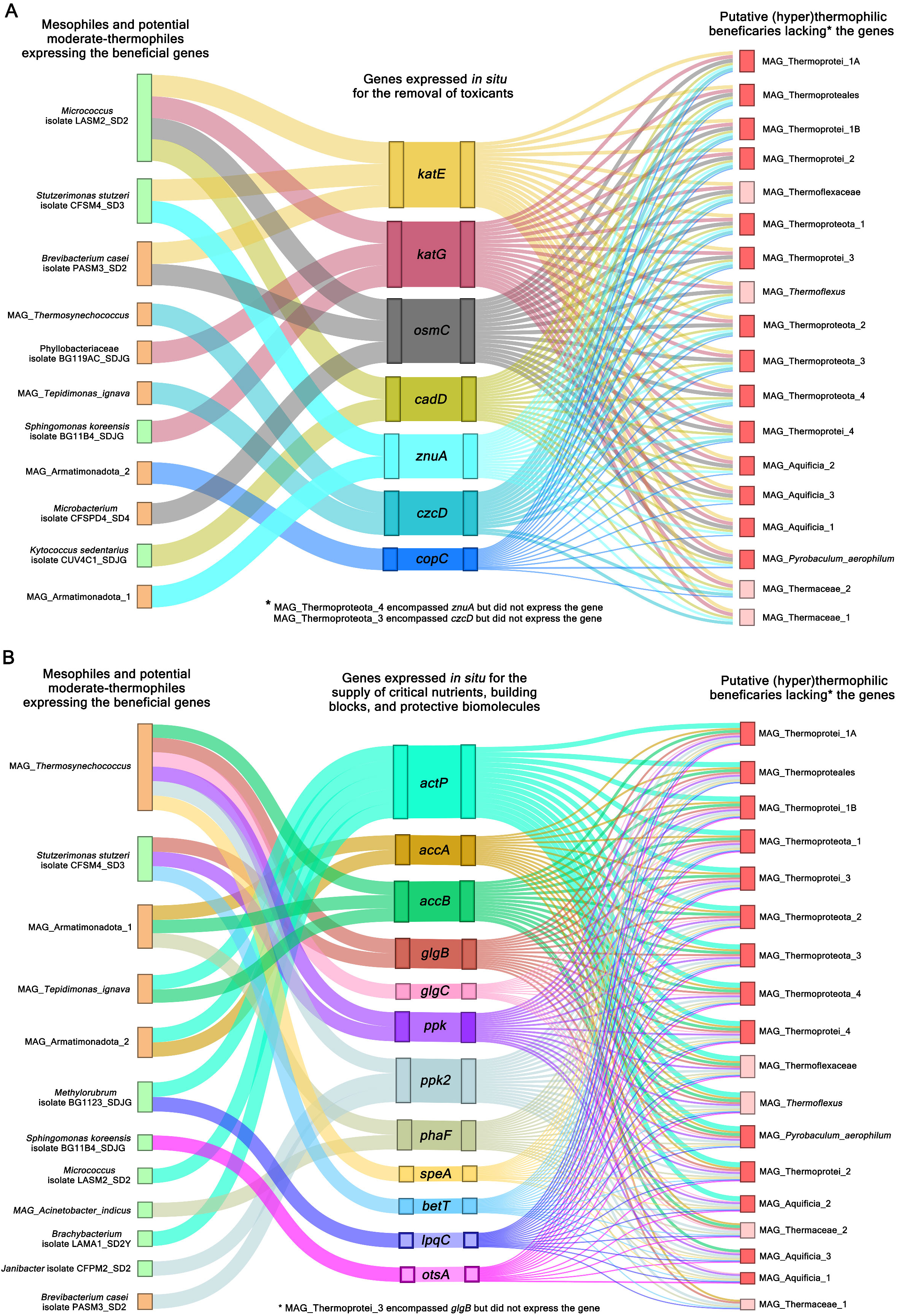
Genes concerned with the (**A**) scavenging of environmental toxicants, and (**B**) supply of nutrients and anti-stress biomolecules, for which metatranscriptomic signatures were detected in one or more mesophile(s) and/or potential moderate-thermophile(s), but not in most of the putative hyperthermophiles and thermophiles, of CMSI vent-water. With regards to the protein products encoded by these genes, and in the context of *in situ* expression, the mesophile(s) and/or potential moderate-thermophile(s) have been envisaged as potential benefactors / suppliers whereas the putative (hyper)thermophiles have been depicted as plausible beneficiaries / recipients. Species names are highlighted by different colors based on their putative thermal affinities (the color code is same as the one used in Fig. 5). EL_gene_ values underlying the information conveyed here are given in Table S78.

The expressed gene-catalogs of a number of CMSI mesophiles, and potential moderate-thermophiles, encompassed indicators (Table S78; Fig. 12B) of their ecological roles as suppliers of critical nutrients and building blocks, in addition to such biomolecules which can protect the geothermal microbiota in general from diverse biophysical stressors^37–39^. For instance, metatranscriptomic reads corresponding to

i. the acetate transporter gene *actP* were detected for three mesophiles and two moderate-thermophiles;
ii. the xylan degradation gene *lpqC* were detected for the mesophilic *Methylorubrum* isolate BG1123_SDJG;
iii. the polyphosphate accumulation genes *ppk* and *ppk2* were detected for two mesophiles and two moderate-thermophiles;
iv. the polyhydroxyalkanoate-granule-formation genes *phaF* were detected for one mesophile and one moderate-thermophile;
v. some or all of the glycogen synthesizing genes *glgA*, *glgB*, *glgC* and *glgP* were detected for one mesophile and three moderate-thermophiles;
vi. some or all of the lipid accumulation genes *accA*, *accB*, *accD*, *fabD*, *fabG*, *fabI*, *plsC*, *plsX*, and/or *plsY* were detected for two mesophiles and seven moderate-thermophiles;
vii. the trehalose producing gene *otsA* were detected for the mesophilic *Sphingomonas* species;
viii. the glycine-betaine transporter gene *betT* were detected for the mesophilic *Stutzerimonas* CFSM4_SD3;
ix. some or all of the polyamine production and transportation genes *speA*, *speB*, *speD*, *speE*, *potA*, and *potD* were detected for one mesophile and four moderate-thermophiles.

### (Hyper)thermophiles as recipients of metabolic services from mesophiles and moderate-thermophiles co-inhabiting the vent-water

The ROS-/metals-detoxification genes such as *osmC*, *katE*, *katG*; *cadD*, and *copC*, or for that matter the nutrient-/biomolecule-producing genes such as *actP*; *lpqC*; *ppk*; *phaF*; *glgB*, *glgC*; *accA*, *accB*, *accD*; *betT*; and *speA* were either not present, or remained unexpressed, in most of the putative hyperthermophiles and thermophiles *in situ* (Table S78). These findings give credibility to the hypothesis that geothermal-vent mesophiles and moderate-thermophiles make critical contributions to ecosystem functioning by scavenging environmental toxicants and supplying vital nutrients and anti-stress biomolecules to the native hyperthermophiles and thermophiles.

Furthermore, corroborative to the above postulate, the *in situ* expressed gene-catalogs of a number of putative hyperthermophiles and thermophiles encompassed key markers of intercellular nanotube formation (Table S78). Such tens of nm wide, hollow, membranous bridges allow microbial cells to connect proximate cells of the same or different species, and in doing so help them to exchange metabolites and genetic information, particularly in the contexts of stress tolerance and biofilm formation^40–42^. Metatranscriptomic reads corresponding to

i. the *ymdB* gene, which encodes a phosphodiesterase essential for accurate nanotube biogenesis and intercellular exchange of molecules, was detected for Aquificia_2, Aquificia_3, and Thermaceae_1;
ii. some or all of the core genes *fliP*, *fliQ* and *fliR*; and *flhA* and *flhB* involved in the construction of nanotubes (as well as flagella) were detected for Aquificia_1, Aquificia_2 and Aquificia_3;
iii. *cdvB*, which is involved in remodeling the archaeal cell envelope to form tubular membrane networks, was detected for most of the CMSI archaea.

These findings suggest that vent-water (hyper)thermophiles may take up different metabolites and cytoplasmic materials from mesophiles and moderate-thermophiles stochastically introduced to the unfamiliar extreme ecosystem. On the flip side of this deduction, *in situ* feasibility of intercellular bridges evokes a new conjecture that molecular aid from (hyper)thermophiles might be a driver of thermal endurance in vent-water mesophiles, over and above the intrinsic^19,43,44^ as well as environment-guided^4,9^ factors already attributed to secondarily acquired thermophilicity.

## Conclusion

This research provides an unprecedented perspective on the habitability, and structure-function dynamics, of geothermal-vent ecosystems. It refutes the notion that vent-epicenters are out of bound territories for microorganisms having no evolutionary background, so no *in vitro* phenotype, of surviving amidst high heat. Instead, the study reveals that the boiling water discharged by a geothermal vent can support a considerably diversified microbiome, which despite being overwhelmingly dominated by (hyper)thermophilic bacteria and archaea, embraces mesophilic immigrants as its regular members. Overall, a geothermal vent is portrayed as an inclusive, multi-community ecosystem, where putative (hyper)thermophiles, potential moderate-thermophiles, and mesophiles live side-by-side, regardless of the large numeral and functional asymmetries existing between, as well as within, these temperature-affinity-based communities. Co-existence of metabolically differentiated, multi-layered microbial communities in a biophysically extreme habitat begets a fundamental paradigm based on which stress-resilient consortia can be optimized for biotechnological applications.

Based on the prevalence:functionality asymmetries recorded across the microbiome, it is inferred that uncoupling of overall metabolism from growth and proliferation may be a general strategy of thermal stress management, irrespective of the adaptational efficacy of the species. This provides a conceptual basis for developing stress-resilient microbial biotechnology platforms where metabolic functionality is preserved independently of growth. Microbial communities that, under extreme biophysical stress, can render active metabolism without growing can be useful in industries where accumulation of biomass is not desired but biocatalysis is necessary.

Pure-culture isolation and consistent metagenome-based detection of a large number of mesophilic and potentially moderately-thermophilic inmates, together with the evidences of their *in situ* transcriptional activity, negate the possibility of the spurious or transitory existence of these bacteria in the explored vent-head. The bulked metatranscriptomic signatures further show that populations of most of the thermally ill-adapted inmates are not only alive, but also active, *in situ*; the gene expression profiles in question illustrate their activities to be aimed essentially at cell-system maintenance and survival, instead of growth and proliferation.

Consistent with conventional expectations, the metatranscriptomic signatures conclude that only those inhabitants of the geothermal vent-head lead a productive and growth-oriented life which are affiliated to (hyper)thermophilic taxa. Counterintuitively, however, the findings envisage the mesophiles and potential moderate-thermophiles as providers of such ancillary metabolic services that can be crucial for the ecological success of the native (hyper)thermophiles. As for the plausible return-services that (hyper)thermophiles may provide to fledgling communities of geothermal-vent mesophiles, only *in vitro* co-culture experiments involving the thermally well-adapted and ill-adapted can tell whether biochemical resources also flow, from the ecologically adept to the inept, under biophysical duress.

## Materials and methods

### Study site and on-field sampling

The Chumathang geothermal area stretches over a few hundred meters along the banks of the river Indus, within the west-central part of the Changthang plateau, in eastern Ladakh. The area harbors several geysers discharging boiling water and fumarole round the year, and a few of them have been studied previously for *in situ* microbiology^21,28^. The vigorous geyser (CMSI) explored here is situated on the bank of the river Indus at latitude 33°2’9.5” N and longitude 78°1’56.6” E (Fig. 1).

Water was sampled from the CMSI vent-epicenter for chemical analysis^17,24^, cell enumeration^19^, pure-culture isolation^16^, and metagenome extraction^9,16,17,19^, as described previously (details given in Supplementary Methods). *In situ* temperature, pH, conductivity, total dissolved solids (TDS), and salinity were recorded using CyberScan PCD 650 Multi-Parameter (Eutech, Thermo Fisher Scientific, USA) following manufacturer’s instructions; the rate of vent-water discharge was also measured *in situ* using soluble tracers as described previously^45^.

### Analytical techniques

Concentrations of boron, cesium, calcium, chromium, cobalt, copper, lithium, magnesium, manganese, potassium, iron, rubidium, strontium, vanadium, and zinc were determined by inductively coupled plasma mass spectrometry, as described previously^9^. Silicon concentration was determined using UV-visible spectrophotometry, whereas chloride, nitrate, sodium, and sulfate were quantified by ion chromatography, as described elsewhere^17^. Total alkalinity was measured by Gran titration method^46^ and dissolved inorganic carbon was measured using a carbon coulometer as described earlier^4^. Dissolved sulfide was estimated from CdS precipitates via methylene blue complex formation, and δ^13^C_DIC_ was measured by isotope ratio mass spectrometry, as described previously^47^.

Using a hemocytometer and an upright fluorescence microscope, density of live, dead, as well as total, microbial cells in a water sample was determined, as described elsewhere^19^, via staining with fluorescein diacetate, propidium iodide, and 4′6-diamidino-2-phenylindole, respectively (details given in Supplementary Methods).

### Isolation of mesophilic bacterial strains: taxonomic identification and draft whole-genome sequencing

Bacterial pure cultures were isolated from the vent-water sample at 37°C, as described previously^16^, using the following culture media: Armbruster medium, BG11 medium, chromogenic culture medium (CHROMagar), EMB Agar, ENDO Agar, MacConkey Agar, Marine Agar 2216, MST medium, PE medium, Pseudomonas medium 2, Sphaerotilus Defined medium, Synthetic Sea Water medium 1, TCBS medium, XLD Agar (detailed procedure given in Supplementary Methods). All the strains isolated being chemoorganoheterotrophic and culturable in Luria broth (LB) were maintained at 37°C in Luria agar slants with an average transfer interval of 20 days. Each strain was classified up to the lowest possible taxonomic rank ascribable on the basis of its 16S rRNA gene sequence identities with known microbial species. For each isolate, genomic DNA was extracted from its stationary-phase LB culture using HiPurA Bacterial Genomic DNA Purification Kit (Himedia, India). The genome was sequenced on an Ion GeneStudio S5 Plus System (Thermo Fisher Scientific, USA), following the manufacturer’s instructions. All Ion S5 reads having Phred score ≥20 were assembled with the help of SPAdes v3.13.0^48^.

### Extraction and sequencing of metagenomes and metatranscriptome

Metagenomic DNA was extracted from the vent-water samples using PureLink Genomic DNA Mini Kit (Thermo Fisher Scientific, USA), as described previously^9,16,17,19^, and detailed in Supplementary Methods. For each of the six individual metagenomic investigations, 30-100 ng DNA was extracted from 10 L vent-water. In tandem with each metagenome extraction procedure, purported kitome DNA was prepared using the same PureLink kit without adding any vent-water sample. Micro-volume spectroscopy using neither NanoDrop nor Qubit technology (both from Thermo Fisher Scientific, USA) detected any DNA in the final eluent of any of the so-called kitomes prepared; therefore, none of the six kitome eluents was subjected to sequencing.

Metatranscriptome (total 132 ng RNA from 50 L vent-water, with RNA integrity number 7.1) was extracted using RNeasy Power Water Kit (Qiagen GmbH, Germany) as detailed in Supplementary Methods. In tandem with the above procedure, purported kitome RNA was prepared using the same RNeasy kit, RNAlater reagent, but no vent-water sample. Micro-volume spectroscopy using neither NanoDrop, nor Qubit or Bioanalyzer, technology detected any RNA in the final eluent of the so-called kitome prepared. The kitome eluent, therefore, was not sequenced subsequently.

Each metagenomic DNA preparation was sequenced for 2×250 nucleotides, paired end reads, using a NovaSeq 6000 System (Illumina Inc., USA). The metatranscriptomic RNA preparation was sequenced for 2×100 nucleotides, paired end reads, using the NovaSeq 6000 System, after removing all rRNA molecules that were present in the stock solution by using Ribo-Zero Gold (Illumina Inc.) as instructed by the manufacturer.

All metagenomic and metatranscriptomic reads were trimmed to remove adapters and filtered for average Phred score ≥20 using Trim Galore v0.6.6 (https://github.com/FelixKrueger/TrimGalore) with default settings. Furthermore, to filter out whatever rRNA-related reads had remained in the metatranscriptomic sequence dataset, the latter was searched against the SILVA_LSURef_138.1_NR99, SILVA_SSURef_138.1_NR99^49^, and rrnDB v5.8^50^, databases using SortMeRNA v4.3.6^51^ in default mode.

### Construction and taxonomic identification of MAGs

Each metagenomic sequence dataset was assembled using Megahit v1.2.9, with minimum contig-length of 300 nucleotides^52^. ≥2500 nucleotide contigs were binned into MAGs using Metabat2, MaxBin2, and CONCOCT^53–55^. MAGs were refined using DASTool, evaluated by CheckM^56,57^, and reported only when they had contamination levels <5%. Taxonomic affiliation of each MAG was determined with the help of the Genome Taxonomy Database Toolkit (GTDB-Tk)^58^ as well as the Type Strain Genome Server (TYGS)^59^; pair-wise comparison of genomes was carried out using Genome-to-Genome Distance Calculator (GGDC) v3.0^60^ and Orthologous Average Nucleotide Identity (OrthoANI)^61^.

### Comprehensive phylogeny of MAGs and genomes

All bacterial and archaeal MAG/genome sequences obtained over the different sampling occasions were curated under two separate datasets, alongside the genome sequences of their closest relatives identified via analyses based on GTDB-Tk and TYGS. The bacterial and archaeal datasets were subjected to phylogenetic clustering of their constituent entities using the up-to-date bacterial core gene set (UBCG v3.0)^62^, and the up-to-date archaeal core gene set (UACG v1.0)^63^, pipelines respectively. For each dataset, a maximum likelihood tree was generated using RAxML^64^, the best substitution model was selected via ModelTest-NG^65^, and the tree was visualized via iTOL^66^.

### Sequence correspondence between the MAGs/genomes and the individual metagenomic datasets

All high-quality reads present in a given metagenomic dataset were mapped, using Bowtie2^67^ in very-sensitive alignment mode, on to the MAG/genome sequences encompassed in the CS-LMGS database. With reference to the mapping experiment in question, the best hit was reported for each read, and the following data were recorded for each MAG/genome included in the CS-LMGS database: (i) number of metagenomic reads matched, (ii) percentage of MAG/genome length matched by metagenomic reads (coverage breadth), (iii) average length of the aligned reads.

### Quantifying *in situ* prevalence of the vent-water species

With reference to a given sampling occasion, Π was calculated as follows.

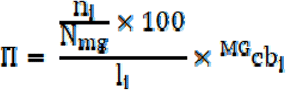

where, n_i_ denotes the number of metagenomic reads matching sequences from the MAG/genome of the i^th^ species; N_mg_ represents the total number of metagenomic reads available for the sampling occasion; l_i_ denotes the length of the i^th^ MAG/genome (in megabases); ^MG^cb_i_ represents the MAG-/genome-wide coverage breadth.

### Annotation of genes within MAGs/genomes

Open reading frames (ORFs), or putative genes, were predicted within the MAG/genome of a CMSI species using Prodigal v2.6.3^68^. The gene-catalog obtained in this way was annotated for protein-coding sequence by searching each ORF against the eggNOG database v5.0^69^ using the online utility eggNOG-mapper v2.1.9^70^ with the default parameters. Furthermore, to identify the COGs to which the different CDSs belonged, the putatively translated form of the Prodigal-derived gene-catalog was annotated by searching against the COG Little Endian version of the Conserved Domain Database using COGclassifier v1.0.5 (https://github.com/moshi4/COGclassifier).

### Identification of gene-sets corresponding to biochemical pathways, structural complexes, and functional units, within a MAG/genome

Defined sets of genes curated in the Kyoto Encyclopedia of Genes and Genomes (KEGG) as coding for specific biochemical pathways, structural complexes, or functional units were identified within a MAG/genome using the web-based tool KEGG Mapper – Reconstruct (https://www.genome.jp/kegg/mapper/reconstruct.html) where the KEGG Orthology identifiers of the CDSs annotated within the MAG/genome were utilized to map the entire CDS-catalog against the information curated in KEGG Modules and BRITE Tables. Because all the MAGs/genomes delineated had partially complete sequences, a module was assumed to be present in a MAG/genome when at least one complete group (block) of genes governing any one step of the concerned pathway was detected in its entirety. In case of modules made of single gene-blocks, their presence within a MAG/genome was ascertained via detection of all the constituent genes of the block in question.

### Sequence correspondence between the MAGs/genomes and the metatranscriptomic dataset

The quality-filtered mRNA-related reads present in the completely-rRNA-read-free metatranscriptomic dataset were mapped, using Bowtie2 in very-sensitive alignment mode, on to the MAG/genome sequences encompassed in the CS-LMGS database. Results of the mapping experiment reported the best hit for each read, and recorded the following data for each MAG/genome included in the CS-LMGS database: (i) number of metatranscriptomic reads matched, (ii) percentage of CDSs matched by metatranscriptomic reads (transcriptional breadth), (iii) average length of the aligned reads.

### Quantifying *in situ* functionality of the vent-water species

For a species included in the CS-LMGS database, coefficient of functionality (Φ), as on 01-Nov-2022, was determined as follows.

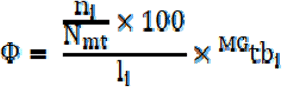

where, n_i_ denotes the number of metatranscriptomic reads matching the MAG/genome of the i^th^ species; N_mt_ represents the total number of metatranscriptomic reads available in the dataset; l_i_ denotes the length of the i^th^ MAG/genome (in megabases); and ^MG^tb_i_ represents the MAG-/genome-wide transcriptional breadth.

### Expression of gene-sets corresponding to pathways, structural complexes, and functional units

For a given MAG/genome, all CDSs expressed *in situ* (CDSs having metatranscriptomic reads mapped on to them) were used to reconstruct KEGG Modules and BRITE Tables, as described above. Subsequently, the *in situ* functional status of a KEGG Module was described as what proportion of its gene-blocks detected completely within the MAG/genome was expressed in entirety (a completed gene-block, in turn, was said to be expressed in its entirety when all its constituent genes were corresponded by metatranscriptomic reads). The functional status of a BRITE Table was described as what proportion of its constituent genes present in the MAG/genome was expressed *in situ* (corresponded by metatranscriptomic reads).

## Acknowledgements

Extensive on-field assistance provided by Sri Asgar Ali, Sri Mohammad Baqir, and Sri Nizam ul Din of Choglamsar, Ladakh, India is gratefully acknowledged. Sri Ayan Dutta provided valuable suggestions regarding extraction of environmental RNA.

## Funding

The study was financed by Bose Institute through Intramural Research Grants. S.D. and J.S. obtained their fellowships from Council of Scientific and Industrial Research, Government of India (GoI). S.C. and M.M. received fellowships from Department of Biotechnology (DBT), GoI. J.G. and S.S. got fellowships from University Grants Commission, GoI. S.P. received fellowship from IODP, NCPOR, Ministry of Earth Sciences (GoI). NM received fellowship from Bose Institute. Bioinformatic analyses were carried out using computational resources available under an EMR project funded by DBT, GoI (BT/PR40174/BTIS/137/45/2022).

## Author contributions

W.G. conceived the study, designed the experiments, interpreted the data, and wrote the paper. S.D. anchored the study, planned and performed the experiments, interpreted and curated the data, and composed the paper. A.P., S.C., J.G., S.P., N.M., M.M., J.S., and S.S. performed the experiments. A.P., A.D., R.C., and A.M. validated the findings and worked towards improving the text and documents. All the authors read and vetted the manuscript.

## Competing interest

The authors declare no competing interest.

## Supplementary data

Supplementary information and data are available online in the form of a Word file named Supplementary_Information.docx, and an Excel file named Supplementary_Dataset.xlsx.

## Data availability

All 16S rRNA gene, and genome, sequences determined for the pure-culture isolates; all metagenome and metatranscriptome sequences determined for the vent-water samples; and sequences of all metagenome-assembled genomes (MAGs) reconstructed, have been deposited to the National Center for Biotechnology Information (NCBI), USA under the BioProject accession number PRJNA1236334.

For every pure-culture isolate, GenBank accession numbers of its 16S rRNA gene sequence and draft whole-genome sequence (WGS) are given below, alongside the Run Accession number of the WGS deposited in the Sequence Read Archive (SRA). *Aeromicrobium* isolate PFPM2_SD2 - PZ321536, JBNFNZ000000000, and SRR33132955; *Agrobacterium pusense* isolate ENURT1_SDJG - PZ321543, JBNFOQ000000000, and SRR33132964; Boseaceae isolate BG11AB_SDJG11 - PZ321541, JBNFOP000000000, and SRR33132963; *Brachybacterium* isolate LAMA1_SD2Y - PZ321545, JBNFOO000000000, and SRR33132952; *Brevibacterium casei* isolate PASM3_SD2 - PZ321546, JBNFOO000000000, and SRR33132948; *Brevibacterium* isolate LAARM2_SD2 - PZ321547, JBNFOO000000000, and SRR33132947; *Brevibacterium* isolate LAARM3_SD1 - PZ321552, JBNFOL000000000, and SRR33132946; *Brevundimonas aurantiaca* isolate CURTG4B_SDJG - PZ321560, JBNFOK000000000, and SRR33132945; *Brevundimonas* isolate CURTG3B2_SDJG - PZ321562, JBNFOJ000000000, and SRR33132944; *Brevundimonas* isolate ENURT2_SDJG - PZ321561, JBNFOI000000000, and SRR33132943; Brucellaceae isolate CFSPD4_SD2 - PZ321565, JBNFOH000000000, and SRR33132942; *Corynebacterium kalidii* isolate LAMA1_SD2W - PZ321568, JBNFOG000000000, and SRR33132962; *Janibacter* isolate CFPM2_SD2 - PZ321569, JBNFOF000000000, and SRR33132961; *Kytococcus sedentarius* isolate CUV4C1_SDJG - PZ321570, JBNFOE000000000, and SRR33132960; *Methylorubrum* isolate BG1123_SDJG - PZ321573, JBNFOD000000000, and SRR33132959; *Microbacterium* isolate CFSPD4_SD4 - PZ321574, JBNFOC000000000, and SRR33132958; *Micrococcus* isolate LASM2_SD2 - PZ321578, JBNFOB000000000, and SRR33132957; *Mycolicibacterium iranicum* isolate BG11O_SDJG - PZ321579, JBNFOA000000000, and SRR33132956; Phyllobacteriaceae isolate BG119AC_SDJG - PZ321583, JBNFNY000000000, and SRR33132954; *Sphingomonas koreensis* isolate BG11B4_SDJG - PZ321630, JBNFNX000000000, and SRR33132953; *Sphingopyxis ummariensis* isolate BG11B2_SDJG - PZ321631, JBNFNW000000000, and SRR33132951; *Stutzerimonas stutzeri* isolate CFSM4_SD3 - PZ321632, JBNFNV000000000, and SRR33132950; *Xanthobacter* isolate BG11Y11_SDJG - PZ321633, JBNFNU000000000, and SRR33132949.

The Run Accession numbers of the six whole-metagenome, and one whole-metatranscriptome, shotgun sequence datasets deposited in the SRA are given below. FN_Nov_21_Mg - SRR32700099, AN_Nov_21_Mg - SRR32700098, FN_Apr_22_Mg - SRR32700097, AN_Apr_22_Mg - SRR32700096, FN_Nov_22_Mg - SRR32700095, AN_Nov_22_Mg - SRR32700094; and WD_Nov_22_Mt - SRR32700093.

GenBank accession numbers of all the 131 MAGs reconstructed in this study have been given in Table S2.

## Notes

### Competing Interest Statement

The authors have declared no competing interest.

### Summary of Updates

As the new title suggests, this new version of the paper is a thorough overhaul in terms of both analyses and interpretations. Specifically the most important addition has been the data on potential ecological role of mesophilic bacteria. All the population-level data have now been re-cast into a community-level higher-order interpretation.

