## Supplementary_Files for "Mesophilic inmates of a geothermal vent-head as providers of critical ecosystem services": Supplementary_Information.docx

**Table of Contents**

**Supplementary Method**

1. On-field sampling
2. Microbial cell density
3. Isolation of mesophilic microbial strains
4. Media used for the isolation of mesophilic bacteria
5. Extraction of metagenomes and metatranscriptome

**Supplementary Results and Discussion**

1. Molecular drivers underlying the asymmetric ecology of putative hyperthermophiles
2. Molecular underpinnings of the numeral/functional laggardness of the ecologically less-efficient archaea
3. Strategies underlying the ecological fitness of the putative thermophiles and potential moderate-thermophiles of CMSI
4. Molecular drivers of the persistence exhibited by the CMSI mesophiles

**Supplementary Tables**

(All the supplementary tables, except S3, have been given in individual sheets of an Excel Workbook named Supplementary_Dataset)

**Table S1.** Physicochemical characteristics of the CMSI vent-water recorded over the six sampling occasions.

**Table S2.** Characteristic features of the bacterial and archaeal population genome bins (MAGs) that were reconstructed from the six different metagenomic datasets, obtained from the CMSI vent-water, over the six sampling occasion.

**Table S3.** Species-level entities discovered in the form of metagenome-assembled genomes (MAGs), over the six different explorations of the CMSI vent-water; maximum temperature (Tmax °C) known for the laboratory growth of the member strains of the lowest-possible taxon to which every MAG-based species could be classified has also been given.

**Table S4.** Pair-wise OrthoANI (cells indicated by pink shades) and dDDH (cells indicated by blue shades) values delineated between the different MAGs and genomes that were retrieved for each bacterial or archaeal species discovered in the CMSI vent-water over the entire exploration period.

**Table S5.** Percentages of reads from the different metagenomic and metatranscriptomic datasets that mapped individually onto the assembled versions of the six different metagenomes.

**Table S6.** Summary of the isolation, identification, and draft genome sequencing of 37°C-growing bacteria from the CMSI vent-water; maximum temperature (Tmax °C) known for the laboratory growth of the member strains of the lowest-possible taxon to which every bacterium isolated could be classified has also been given.

**Table S7.** MAGs and genomes included in the CS-LMGS database as representatives of the 66 bacterial and archaeal species discovered in the CMSI vent-water over the entire exploration period.

**Table S8.** Delineation of coefficients of prevalence (Π) for each species included in the CS-LMGS database, over the six different sampling occasions, on the basis of the sequence-correspondence levels recorded between the concerned MAG/genome and the individual metagenomic datasets; coefficients of functionality (Φ) have also been delineated for each species included in the CS-LMGS database, on the basis of the sequence-correspondence levels recorded between the concerned MAG/genome and the sole metatranscriptomic dataset retrieved through the forenoon and afternoon of 01-November-2022.

**Table S9.** Coding sequence (CDS) catalog of MAG_Aquificia_2 obtained by annotating its gene-catalog with the help of COGclassifier and eggNOG-mapper; for each gene/CDS identified, the number of metatranscriptomic reads matching its sequence have been given alongside the expression level of the gene/CDS (EL_gene_).

**Table S10.** Coding sequence (CDS) catalog of MAG_Thermoprotei_1A obtained by annotating its gene-catalog with the help of COGclassifier and eggNOG-mapper; for each gene/CDS identified, the number of metatranscriptomic reads matching its sequence have been given alongside the expression level of the gene/CDS (EL_gene_).

**Table S11.** Coding sequence (CDS) catalog of MAG_Thermoproteales obtained by annotating its gene-catalog with the help of COGclassifier and eggNOG-mapper; for each gene/CDS identified, the number of metatranscriptomic reads matching its sequence have been given alongside the expression level of the gene/CDS (EL_gene_).

**Table S12.** Coding sequence (CDS) catalog of MAG_Thermoprotei_1B obtained by annotating its gene-catalog with the help of COGclassifier and eggNOG-mapper; for each gene/CDS identified, the number of metatranscriptomic reads matching its sequence have been given alongside the expression level of the gene/CDS (EL_gene_).

**Table S13.** Coding sequence (CDS) catalog of MAG_Aquificia_3 obtained by annotating its gene-catalog with the help of COGclassifier and eggNOG-mapper; for each gene/CDS identified, the number of metatranscriptomic reads matching its sequence have been given alongside the expression level of the gene/CDS (EL_gene_).

**Table S14.** Coding sequence (CDS) catalog of MAG_Aquificia_1 obtained by annotating its gene-catalog with the help of COGclassifier and eggNOG-mapper; for each gene/CDS identified, the number of metatranscriptomic reads matching its sequence have been given alongside the expression level of the gene/CDS (EL_gene_).

**Table S15.** Coding sequence (CDS) catalog of MAG_Thermaceae_2 obtained by annotating its gene-catalog with the help of COGclassifier and eggNOG-mapper; for each gene/CDS identified, the number of metatranscriptomic reads matching its sequence have been given alongside the expression level of the gene/CDS (EL_gene_).

**Table S16.** Coding sequence (CDS) catalog of MAG_Thermaceae_1 obtained by annotating its gene-catalog with the help of COGclassifier and eggNOG-mapper; for each gene/CDS identified, the number of metatranscriptomic reads matching its sequence have been given alongside the expression level of the gene/CDS (EL_gene_).

**Table S17.** Coding sequence (CDS) catalog of MAG_Armatimonadota_1 obtained by annotating its gene-catalog with the help of COGclassifier and eggNOG-mapper; for each gene/CDS identified, the number of metatranscriptomic reads matching its sequence have been given alongside the expression level of the gene/CDS (EL_gene_).

**Table S18.** Coding sequence (CDS) catalog of MAG_Bacteria_1 obtained by annotating its gene-catalog with the help of COGclassifier and eggNOG-mapper; for each gene/CDS identified, the number of metatranscriptomic reads matching its sequence have been given alongside the expression level of the gene/CDS (EL_gene_).

**Table S19.** Coding sequence (CDS) catalog of MAG_*Thermosynechococcus* obtained by annotating its gene-catalog with the help of COGclassifier and eggNOG-mapper; for each gene/CDS identified, the number of metatranscriptomic reads matching its sequence have been given alongside the expression level of the gene/CDS (EL_gene_).

**Table S20.** Coding sequence (CDS) catalog of MAG_Thermoprotei_2 obtained by annotating its gene-catalog with the help of COGclassifier and eggNOG-mapper; for each gene/CDS identified, the number of metatranscriptomic reads matching its sequence have been given alongside the expression level of the gene/CDS (EL_gene_).

**Table S21.** Coding sequence (CDS) catalog of MAG_Thermoflexaceae obtained by annotating its gene-catalog with the help of COGclassifier and eggNOG-mapper; for each gene/CDS identified, the number of metatranscriptomic reads matching its sequence have been given alongside the expression level of the gene/CDS (EL_gene_).

**Table S22.** Coding sequence (CDS) catalog of MAG_Thermoproteota_1 obtained by annotating its gene-catalog with the help of COGclassifier and eggNOG-mapper; for each gene/CDS identified, the number of metatranscriptomic reads matching its sequence have been given alongside the expression level of the gene/CDS (EL_gene_).

**Table S23.** Coding sequence (CDS) catalog of MAG_Thermoprotei_3 obtained by annotating its gene-catalog with the help of COGclassifier and eggNOG-mapper; for each gene/CDS identified, the number of metatranscriptomic reads matching its sequence have been given alongside the expression level of the gene/CDS (EL_gene_).

**Table S24.** Coding sequence (CDS) catalog of MAG_Halomonadaceae obtained by annotating its gene-catalog with the help of COGclassifier and eggNOG-mapper; for each gene/CDS identified, the number of metatranscriptomic reads matching its sequence have been given alongside the expression level of the gene/CDS (EL_gene_).

**Table S25.** Coding sequence (CDS) catalog of MAG_*Pyrobaculum*_*aerophilum* obtained by annotating its gene-catalog with the help of COGclassifier and eggNOG-mapper; for each gene/CDS identified, the number of metatranscriptomic reads matching its sequence have been given alongside the expression level of the gene/CDS (EL_gene_).

**Table S26.** Coding sequence (CDS) catalog of MAG_*Vibrio*_*metschnikovii* obtained by annotating its gene-catalog with the help of COGclassifier and eggNOG-mapper; for each gene/CDS identified, the number of metatranscriptomic reads matching its sequence have been given alongside the expression level of the gene/CDS (EL_gene_).

**Table S27.** Coding sequence (CDS) catalog of MAG_Armatimonadota_2 obtained by annotating its gene-catalog with the help of COGclassifier and eggNOG-mapper; for each gene/CDS identified, the number of metatranscriptomic reads matching its sequence have been given alongside the expression level of the gene/CDS (EL_gene_).

**Table S28.** Coding sequence (CDS) catalog of MAG_*Thermoflexus* obtained by annotating its gene-catalog with the help of COGclassifier and eggNOG-mapper; for each gene/CDS identified, the number of metatranscriptomic reads matching its sequence have been given alongside the expression level of the gene/CDS (EL_gene_).

**Table S29.** Coding sequence (CDS) catalog of MAG_Thermoproteota_2 obtained by annotating its gene-catalog with the help of COGclassifier and eggNOG-mapper; for each gene/CDS identified, the number of metatranscriptomic reads matching its sequence have been given alongside the expression level of the gene/CDS (EL_gene_).

**Table S30.** Coding sequence (CDS) catalog of MAG_Bacteria_3 obtained by annotating its gene-catalog with the help of COGclassifier and eggNOG-mapper; for each gene/CDS identified, the number of metatranscriptomic reads matching its sequence have been given alongside the expression level of the gene/CDS (EL_gene_).

**Table S31.** Coding sequence (CDS) catalog of MAG_Gammaproteobacteria obtained by annotating its gene-catalog with the help of COGclassifier and eggNOG-mapper; for each gene/CDS identified, the number of metatranscriptomic reads matching its sequence have been given alongside the expression level of the gene/CDS (EL_gene_).

**Table S32.** Coding sequence (CDS) catalog of MAG_Bacteria_4 obtained by annotating its gene-catalog with the help of COGclassifier and eggNOG-mapper; for each gene/CDS identified, the number of metatranscriptomic reads matching its sequence have been given alongside the expression level of the gene/CDS (EL_gene_).

**Table S33.** Coding sequence (CDS) catalog of MAG_*Tepidimonas*_*ignava* obtained by annotating its gene-catalog with the help of COGclassifier and eggNOG-mapper; for each gene/CDS identified, the number of metatranscriptomic reads matching its sequence have been given alongside the expression level of the gene/CDS (EL_gene_).

**Table S34.** Coding sequence (CDS) catalog of MAG_Thermoproteota_3 obtained by annotating its gene-catalog with the help of COGclassifier and eggNOG-mapper; for each gene/CDS identified, the number of metatranscriptomic reads matching its sequence have been given alongside the expression level of the gene/CDS (EL_gene_).

**Table S35.** Coding sequence (CDS) catalog of MAG_Thermoproteota_4 obtained by annotating its gene-catalog with the help of COGclassifier and eggNOG-mapper; for each gene/CDS identified, the number of metatranscriptomic reads matching its sequence have been given alongside the expression level of the gene/CDS (EL_gene_).

**Table S36.** Coding sequence (CDS) catalog of *Stutzerimonas* *stutzeri* isolate CFSM4_SD3 obtained by annotating its gene-catalog with the help of COGclassifier and eggNOG-mapper; for each gene/CDS identified, the number of metatranscriptomic reads matching its sequence have been given alongside the expression level of the gene/CDS (EL_gene_).

**Table S37.** Coding sequence (CDS) catalog of *Agrobacterium* *pusense* isolate ENURT1_SDJG obtained by annotating its gene-catalog with the help of COGclassifier and eggNOG-mapper; for each gene/CDS identified, the number of metatranscriptomic reads matching its sequence have been given alongside the expression level of the gene/CDS (EL_gene_).

**Table S38.** Coding sequence (CDS) catalog of MAG_Thermoprotei_4 MAG_Thermoproteales obtained by annotating its gene-catalog with the help of COGclassifier and eggNOG-mapper; for each gene/CDS identified, the number of metatranscriptomic reads matching its sequence have been given alongside the expression level of the gene/CDS (EL_gene_).

**Table S39.** Coding sequence (CDS) catalog of MAG_*Sphingomonas* obtained by annotating its gene-catalog with the help of COGclassifier and eggNOG-mapper; for each gene/CDS identified, the number of metatranscriptomic reads matching its sequence have been given alongside the expression level of the gene/CDS (EL_gene_).

**Table S40.** Coding sequence (CDS) catalog of MAG_Bacteria_2 obtained by annotating its gene-catalog with the help of COGclassifier and eggNOG-mapper; for each gene/CDS identified, the number of metatranscriptomic reads matching its sequence have been given alongside the expression level of the gene/CDS (EL_gene_).

**Table S41.** Coding sequence (CDS) catalog of *Brevundimonas* isolate ENURT2_SDJG obtained by annotating its gene-catalog with the help of COGclassifier and eggNOG-mapper; for each gene/CDS identified, the number of metatranscriptomic reads matching its sequence have been given alongside the expression level of the gene/CDS (EL_gene_).

**Table S42.** Coding sequence (CDS) catalog of *Methylorubrum* isolate BG1123_SDJG obtained by annotating its gene-catalog with the help of COGclassifier and eggNOG-mapper; for each gene/CDS identified, the number of metatranscriptomic reads matching its sequence have been given alongside the expression level of the gene/CDS (EL_gene_).

**Table S43.** Coding sequence (CDS) catalog of MAG_Boseaceae obtained by annotating its gene-catalog with the help of COGclassifier and eggNOG-mapper; for each gene/CDS identified, the number of metatranscriptomic reads matching its sequence have been given alongside the expression level of the gene/CDS (EL_gene_).

**Table S44.** Coding sequence (CDS) catalog of MAG_*Brevundimonas*_1 obtained by annotating its gene-catalog with the help of COGclassifier and eggNOG-mapper; for each gene/CDS identified, the number of metatranscriptomic reads matching its sequence have been given alongside the expression level of the gene/CDS (EL_gene_).

**Table S45.** Coding sequence (CDS) catalog of *Brevibacterium* *casei* isolate PASM3_SD2 obtained by annotating its gene-catalog with the help of COGclassifier and eggNOG-mapper; for each gene/CDS identified, the number of metatranscriptomic reads matching its sequence have been given alongside the expression level of the gene/CDS (EL_gene_).

**Table S46.** Coding sequence (CDS) catalog of MAG_*Psychrobacter* obtained by annotating its gene-catalog with the help of COGclassifier and eggNOG-mapper; for each gene/CDS identified, the number of metatranscriptomic reads matching its sequence have been given alongside the expression level of the gene/CDS (EL_gene_).

**Table S47.** Coding sequence (CDS) catalog of *Brevibacterium* isolate LAARM3_SD1 obtained by annotating its gene-catalog with the help of COGclassifier and eggNOG-mapper; for each gene/CDS identified, the number of metatranscriptomic reads matching its sequence have been given alongside the expression level of the gene/CDS (EL_gene_).

**Table S48.** Coding sequence (CDS) catalog of MAG_*Brevundimonas*_3 obtained by annotating its gene-catalog with the help of COGclassifier and eggNOG-mapper; for each gene/CDS identified, the number of metatranscriptomic reads matching its sequence have been given alongside the expression level of the gene/CDS (EL_gene_).

**Table S49.** Coding sequence (CDS) catalog of *Microbacterium* isolate CFSPD4_SD4 obtained by annotating its gene-catalog with the help of COGclassifier and eggNOG-mapper; for each gene/CDS identified, the number of metatranscriptomic reads matching its sequence have been given alongside the expression level of the gene/CDS (EL_gene_).

**Table S50.** Coding sequence (CDS) catalog of *Micrococcus* isolate LASM2_SD2 obtained by annotating its gene-catalog with the help of COGclassifier and eggNOG-mapper; for each gene/CDS identified, the number of metatranscriptomic reads matching its sequence have been given alongside the expression level of the gene/CDS (EL_gene_).

**Table S51.** Coding sequence (CDS) catalog of MAG_*Georgenia* obtained by annotating its gene-catalog with the help of COGclassifier and eggNOG-mapper; for each gene/CDS identified, the number of metatranscriptomic reads matching its sequence have been given alongside the expression level of the gene/CDS (EL_gene_).

**Table S52.** Coding sequence (CDS) catalog of *Brachybacterium* isolate LAMA1_SD2Y obtained by annotating its gene-catalog with the help of COGclassifier and eggNOG-mapper; for each gene/CDS identified, the number of metatranscriptomic reads matching its sequence have been given alongside the expression level of the gene/CDS (EL_gene_).

**Table S53.** Coding sequence (CDS) catalog of Brucellaceae isolate CFSPD4_SD2 obtained by annotating its gene-catalog with the help of COGclassifier and eggNOG-mapper; for each gene/CDS identified, the number of metatranscriptomic reads matching its sequence have been given alongside the expression level of the gene/CDS (EL_gene_).

**Table S54.** Coding sequence (CDS) catalog of MAG_*Acinetobacter*_*indicus* obtained by annotating its gene-catalog with the help of COGclassifier and eggNOG-mapper; for each gene/CDS identified, the number of metatranscriptomic reads matching its sequence have been given alongside the expression level of the gene/CDS (EL_gene_).

**Table S55.** Coding sequence (CDS) catalog of *Kytococcus* *sedentarius* isolate CUV4C1_SDJG obtained by annotating its gene-catalog with the help of COGclassifier and eggNOG-mapper; for each gene/CDS identified, the number of metatranscriptomic reads matching its sequence have been given alongside the expression level of the gene/CDS (EL_gene_).

**Table S56.** Coding sequence (CDS) catalog of *Brevundimonas* *aurantiaca* isolate CURTG4B_SDJG obtained by annotating its gene-catalog with the help of COGclassifier and eggNOG-mapper; for each gene/CDS identified, the number of metatranscriptomic reads matching its sequence have been given alongside the expression level of the gene/CDS (EL_gene_).

**Table S57.** Coding sequence (CDS) catalog of *Janibacter* isolate CFPM2_SD2 obtained by annotating its gene-catalog with the help of COGclassifier and eggNOG-mapper; for each gene/CDS identified, the number of metatranscriptomic reads matching its sequence have been given alongside the expression level of the gene/CDS (EL_gene_).

**Table S58.** Coding sequence (CDS) catalog of MAG_Burkholderiales_2 obtained by annotating its gene-catalog with the help of COGclassifier and eggNOG-mapper; for each gene/CDS identified, the number of metatranscriptomic reads matching its sequence have been given alongside the expression level of the gene/CDS (EL_gene_).

**Table S59.** Coding sequence (CDS) catalog of MAG_Burkholderiales_1 obtained by annotating its gene-catalog with the help of COGclassifier and eggNOG-mapper; for each gene/CDS identified, the number of metatranscriptomic reads matching its sequence have been given alongside the expression level of the gene/CDS (EL_gene_).

**Table S60.** Coding sequence (CDS) catalog of Boseaceae isolate BG11AB_SDJG11 obtained by annotating its gene-catalog with the help of COGclassifier and eggNOG-mapper; for each gene/CDS identified, the number of metatranscriptomic reads matching its sequence have been given alongside the expression level of the gene/CDS (EL_gene_).

**Table S61.** Coding sequence (CDS) catalog of *Aeromicrobium* isolate PFPM2_SD2 obtained by annotating its gene-catalog with the help of COGclassifier and eggNOG-mapper; for each gene/CDS identified, the number of metatranscriptomic reads matching its sequence have been given alongside the expression level of the gene/CDS (EL_gene_).

**Table S62.** Coding sequence (CDS) catalog of Phyllobacteriaceae isolate BG119AC_SDJG obtained by annotating its gene-catalog with the help of COGclassifier and eggNOG-mapper; for each gene/CDS identified, the number of metatranscriptomic reads matching its sequence have been given alongside the expression level of the gene/CDS (EL_gene_).

**Table S63.** Coding sequence (CDS) catalog of *Xanthobacter* isolate BG11Y11_SDJG obtained by annotating its gene-catalog with the help of COGclassifier and eggNOG-mapper; for each gene/CDS identified, the number of metatranscriptomic reads matching its sequence have been given alongside the expression level of the gene/CDS (EL_gene_).

**Table S64.** Coding sequence (CDS) catalog of *Mycolicibacterium* *iranicum* isolate BG11O_SDJG obtained by annotating its gene-catalog with the help of COGclassifier and eggNOG-mapper; for each gene/CDS identified, the number of metatranscriptomic reads matching its sequence have been given alongside the expression level of the gene/CDS (EL_gene_).

**Table S65.** Coding sequence (CDS) catalog of *Corynebacterium* *kalidii* isolate LAMA1_SD2W obtained by annotating its gene-catalog with the help of COGclassifier and eggNOG-mapper; for each gene/CDS identified, the number of metatranscriptomic reads matching its sequence have been given alongside the expression level of the gene/CDS (EL_gene_).

**Table S66.** Coding sequence (CDS) catalog of MAG_Sphingomonadales obtained by annotating its gene-catalog with the help of COGclassifier and eggNOG-mapper; for each gene/CDS identified, the number of metatranscriptomic reads matching its sequence have been given alongside the expression level of the gene/CDS (EL_gene_).

**Table S67.** Coding sequence (CDS) catalog of MAG_Cyanobacteriota_1 obtained by annotating its gene-catalog with the help of COGclassifier and eggNOG-mapper; for each gene/CDS identified, the number of metatranscriptomic reads matching its sequence have been given alongside the expression level of the gene/CDS (EL_gene_).

**Table S68.** Coding sequence (CDS) catalog of MAG_Cyanobacteriota_2 obtained by annotating its gene-catalog with the help of COGclassifier and eggNOG-mapper; for each gene/CDS identified, the number of metatranscriptomic reads matching its sequence have been given alongside the expression level of the gene/CDS (EL_gene_).

**Table S69.** Coding sequence (CDS) catalog of *Brevundimonas* isolate CURTG3B2_SDJG obtained by annotating its gene-catalog with the help of COGclassifier and eggNOG-mapper; for each gene/CDS identified, the number of metatranscriptomic reads matching its sequence have been given alongside the expression level of the gene/CDS (EL_gene_).

**Table S70.** Coding sequence (CDS) catalog of *Sphingomonas* *koreensis* isolate BG11B4_SDJG obtained by annotating its gene-catalog with the help of COGclassifier and eggNOG-mapper; for each gene/CDS identified, the number of metatranscriptomic reads matching its sequence have been given alongside the expression level of the gene/CDS (EL_gene_).

**Table S71.** Coding sequence (CDS) catalog of *Sphingopyxis* *ummariensis* isolate BG11B2_SDJG obtained by annotating its gene-catalog with the help of COGclassifier and eggNOG-mapper; for each gene/CDS identified, the number of metatranscriptomic reads matching its sequence have been given alongside the expression level of the gene/CDS (EL_gene_).

**Table S72.** Coding sequence (CDS) catalog of MAG_Burkholderiales_3 obtained by annotating its gene-catalog with the help of COGclassifier and eggNOG-mapper; for each gene/CDS identified, the number of metatranscriptomic reads matching its sequence have been given alongside the expression level of the gene/CDS (EL_gene_).

**Table S73.** Coding sequence (CDS) catalog of MAG_Moraxellaceae obtained by annotating its gene-catalog with the help of COGclassifier and eggNOG-mapper; for each gene/CDS identified, the number of metatranscriptomic reads matching its sequence have been given alongside the expression level of the gene/CDS (EL_gene_).

**Table S74.** Coding sequence (CDS) catalog of *Brevibacterium* isolate LAARM2_SD2 obtained by annotating its gene-catalog with the help of COGclassifier and eggNOG-mapper; for each gene/CDS identified, the number of metatranscriptomic reads matching its sequence have been given alongside the expression level of the gene/CDS (EL_gene_).

**Table S75.** Delineation of the expression levels of the different COG-categories (EL_CC_ values) in individual CMSI species; data underlying every EL_CC_ calculation, namely the median EL_gene_ and ^CC^tb values determined for each COG-category expressed by a species, have also been shown.

**Table S76.** Metatranscriptomic signatures recorded for the *in situ* expression of different biochemical pathways, structural complexes, and functional units (KEGG Modules) by the 66 CMSI vent-water species.

**Table S77.** Chemolithotrophy- and autotrophy-related genes expressed by CMSI vent-water species, which incidentally were all putative (hyper)thermophiles.

**Table S78.** Ecologically significant genes expressed by one or more mesophile(s) and/or potential moderate thermophile(s) of the CMSI vent-water.

**Table S79.** Correlations delineated between the Φ or Π_μ_ values of the 66 CMSI species on one hand and their EL_CC_ values for the individual COG-categories on the other.

**Supplementary References**

1. References used in Table S3
2. References used in Table S6
3. References used in Supplementary Methods
4. References used in Supplementary Results and Discussion

**Supplementary Figures**

**Figure S1.** Alignment statistics underlying the results obtained for the metagenomic and metatranscriptomic read mapping experiments conducted upon the CS-LMGS database.

**Figure S2.** Scatter plot showing the Φ of each CMSI species as a function of its Π_μ_.

**Supplementary Methods**

**On-field sampling**

For the quantification of metallic elements, 100 mL vent-water was collected via addition of 400 µL HNO_3_ (69% w/v, pH ≤2). For the quantification of non-metallic elements, as well as for measuring total alkalinity (TA) and dissolved inorganic carbon (DIC), another 100 mL water sample was collected via filtration through a 0.22 µm mixed cellulose ester membrane (Merck Life Science Private Limited, India) to stop all biotic activity within the sample. For the quantification of dissolved sulfides water samples were precipitated with 2 M Cd(NO_3_)_2_ as described previously (Roy et al., 2020a).

For the estimation of microbial cell density (Mondal et al., 2024), 1 L vent-water was filtered through a single sterile 0.22 μm mixed cellulose ester membrane, which was then rehydrated and preserved in a solution of 0.9% NaCl in 15% glycerol (glycerol-saline), wrapped with self-sealing thermoplastic films (Tarsons Products Limited, India), and transported to the laboratory under ambient temperature conditions.

For the isolation of pure cultures (Roy et al., 2016), total 10 L vent-water was filtered in 10 different 1 L batches, through 10 individual sterile 0.22 μm mixed cellulose ester membranes. Each membrane was rehydrated and preserved in 5 mL glycerol-saline solution contained in a 10 mL sterile cryovial. All the 10 cryovials lined up in this way were wrapped with self-sealing thermoplastic films, and transported to the laboratory under ambient temperature conditions. Additionally, 1 L unfiltered vent-water was collected in sterile borosilicate glass bottles for the same purpose.

For metagenome extraction (Roy et al., 2020b; Mondal et al., 2022a, 2024), microbial cells were filtered out from a total 10 L of vent-water, using a suite of 10 sterile 0.22 μm mixed cellulose ester membranes. 1 L vent-water was passed through each membrane, following which the membrane was folded and immersed in 5 mL sterile 50 mM Tris:EDTA (pH 7.8) contained in a 10 mL cryovial. All the 10 membrane-containing cryovials lined up in this way were wrapped with self-sealing thermoplastic films, and transported under refrigeration to the laboratory.

For the extraction of metatranscriptome, microbial cells were filtered out from a total 50 L vent-water using a series of 50 sterile 0.22 μm mixed cellulose ester membranes. 1 L vent-water was passed through each membrane, following which the membrane was folded and immersed immediately in 5 mL RNAlater (Sigma-Aldrich, Merck KGaA, Germany) contained in 10 mL sterile cryovials. All the 50 cryovials lined up in this way were wrapped with self-sealing thermoplastic films, and transported under refrigeration to the laboratory.

**Microbial cell density**

On a given sampling occasion, total microbial cell density was determined for the CMSI vent-water, alongside the density of metabolically active as well as inactive cells in the same water sample. All the microbial cells that had been filtered out on field from 1 L vent-water were first dislodged from the 0.22 μm mixed cellulose ester membrane by shredding the membrane with sterile scissors, and then vortexing for 30 min, within the same glycerol-saline containing cryovial in which the membrane had been placed on field. The vial was centrifuged at 1000 *g* for 5 seconds to allow the filter shreds to settle at the bottom of the vial. The supernatant was then collected without disturbing the filter debris lying at the bottom of the vial. The volume of the suspension was measured precisely, and then used for microscopic analyses with the supposition that it contained all the microbial cells which were there in the 1 L vent-water which had been passed through the membrane on field.

Three 50 μL aliquots of the cell suspension were taken and each of them was mixed with 4 μL of a 4′6-diamidino-2-phenylindole (DAPI) solution that had a concentration of 10 µg mL^-1^. All the three mixtures were kept in the dark, at 37°C, for 15 minutes (Dong et al., 2016). After incubation, each of the three stained cell populations was washed twice with sterile phosphate buffered saline (PBS) solution and re-suspended in 50 µL PBS. Finally, 20 µL of each stained and washed cell suspension was dispensed on to a hemocytometer (Paul Marienfeld GmbH & Co. KG, Germany) and examined under an upright fluorescence microscope (Olympus BX53 Digital, Olympus Corporation, Japan) following standard procedure described previously (Quan et al., 2015; Mondal et al., 2024). Finally, the total microbial cell density of the vent-water sample was reported as the average of the three independent data that were obtained from the three hemocytometric experiments carried out as per the guidelines published elsewhere (Absher 1973).

Three 50 μL aliquots of the cell suspension were taken and each of them was mixed with 2 μL of a fluorescein diacetate (FDA) solution that had a concentration of 25 mg mL^-1^. All the three mixtures were kept in the dark, at 37°C, for 15 minutes (Jones and Senft, 1985). After incubation, each of the three stained cell populations was washed, and analyzed using a hemocytometer and the upright fluorescence microscope as stated above. Finally, the density of metabolically active cells in the vent-water sample was reported as the average of the three independent data that were obtained from the three hemocytometric experiments.

Three 50 μL aliquots of the cell suspension were taken and each of them was mixed with 2 μL of a propidium iodide (PI) solution that had a concentration of 5 mg mL^-1^. All the three mixtures were kept in the dark, at 37°C, for 15 minutes (Jones and Senft, 1985), following which each of the three stained cell populations was washed, and analyzed using a hemocytometer and the upright fluorescence microscope as stated above. Eventually, the density of metabolically inactive cells was reported as the average of the three independent data obtained from the three hemocytometric experiments.

**Isolation of mesophilic microbial strains**

The 0.22 μm mixed cellulose ester membranes which were earmarked for pure-culture isolation and preserved in a solution of 0.9% NaCl in 15% glycerol (glycerol-saline) were shredded with sterile scissors inside the vials in which they were brought from the field. All the 10 vials that were in hand for this purpose were vortexed for 15 minutes, and the filter-shreds were allowed to settle down; the supernatants were then collected in a fresh sterilized vial without disturbing the debris. The supernatants collected from the 10 individual vials were pooled to get an approximately 50 mL cell suspension in glycerol-saline solution. This final suspension was used for individual spread-plating experiments carried out at 37°C on the following solidified culture media: Armbruster medium, BG11 medium, chromogenic culture medium (CHROMagar), EMB Agar, ENDO Agar, MacConkey Agar, Marine Agar 2216, MST medium, PE medium, Pseudomonas medium 2, Sphaerotilus Defined medium, Synthetic Sea Water medium 1, TCBS medium, XLD Agar. Additionally, 100 mL unfiltered vent-water was incubated at 37°C for 30 days, after which aliquots of the water were spread on to agar plates of the aforesaid media types. From the microbial lawns appearing on the different agar plates, pure-culture strains were isolated, via dilution streaking, as visually distinct single colonies.

**Media used for the isolation of mesophilic bacteria**

**Armbruster medium** (Chernousova et al., 2009)

Armbruster medium contained (L^-1^ distilled water): 100mg sodium lactate, 1.7mg NH_4_Cl, 8.5mg KH_2_PO_4_, 21.5mg K_2_HPO_4_, 34.4mg Na_2_HPO_4_.7H_2_O, 22.5mg MgSO_4_.7H_2_O, 27.5mg CaCl_2_, 0.25mg FeCl_3_.6H_2_O.

**BG11 medium** (Atlas 2010)

BG11 medium contained (L^-1^ distilled water): 1.5g NaNO_3_, 0.075g MgSO_4_·7H_2_O, 0.04g K_2_HPO_4_, 0.036g CaCl_2_·2H_2_O, 0.02g Na_2_CO_3_, 6.0mg citric acid, 6.0mg ferric ammonium citrate, 10.0g agar, 1.0mg disodium EDTA, 1.0mL trace metal mix A5.

**CHROMagar** (Atlas 2010)

CHROMagar contained (L^-1^ distilled water): 16.0g peptone, 16.0g meat extract, 16.0g yeast extract, 15.0g agar, 2.0g chromogenic mix.

**EMB Agar** (Atlas 2010)

10.0g pancreatic digest of casein, 5.0g lactose, 5.0g sucrose, 2.0g K_2_HPO_4_, 0.4g eosin Y, 0.065g methylene blue, 13.5g agar.

**ENDO Agar** (Atlas 2010)

ENDO Agar contained (L^-1^ distilled water): 10.0g peptic digest of animal tissue, 10.0g lactose, 3.5g K_2_HPO_4_, 2.5g Na_2_SO_3_, 0.5g basic fuchsin, 15.0g agar.

**MacConkey Agar** (Atlas 2010)

MacConkey Agar contained (L^-1^ distilled water): 20.0g peptone, 10.0g lactose, 5.0g bile salts, 5.0g NaCl, 0.075g neutral red, 12.0g agar.

**Marine Agar 2216** (Atlas 2010)

Marine Agar 2216 contained (L^-1^ distilled water): 19.45g NaCl, 8.8g MgCl_2_, 5.0g peptone, 3.24g Na_2_SO_3_, 1.8g CaCl_2_, 1.0g yeast extract, 0.55g KCl, 0.16g NaHCO_3_, 0.1g ferric citrate, 0.08g KBr, 0.03g SrCl_2_, 0.02g H_3_BO_3_, 8.0mg Na_2_HPO_4_, 4.0mg Na_2_SiO_3_, 2.4mg NaF, 1.6mg NH_4_NO_3_, 15.0g agar.

**MST medium** (Ghosh and Roy, 2006)

MST medium contained (L^-1^ distilled water): 1g NH_4_Cl, 4g K_2_HPO_4_, 1·5g KH_2_PO_4_, 0·5g MgSO_4_.7H_2_O, and 5·0ml trace metals solution, 20mM Na_2_S_2_O_3_.5H_2_O supplemented with vitamin mixture (10mg each of nicotinic acid, pantothenic acid, pyridoxine, thiamin, p-aminobenzoic acid, riboflavin, and biotin L^−1^).

**PE medium** (Hirose et al., 2020)

PE medium contained (L^-1^ distilled water): 0.5g sodium glutamate, 0.5g sodium succinate, 0.5g sodium acetate, 0.5g casamino acids, 0.5g sodium thiosulfate, 0.5g ammonium sulfate, 1ml of vitamin mixture, 5 ml of 1M phosphate buffer, and 5 ml of a basal salt solution.

**Pseudomonas medium 2** (Atlas 2010)

Pseudomonas medium 2 contained (L^-1^ distilled water): 6.0g Na_2_HPO_4_·12H_2_O, 5.0g succinic acid, 2.4g KH_2_PO_4_, 1.0g NH_4_Cl, 0.5g MgSO_4_·7H_2_O, 0.01g CaCl_2_·6H_2_O, 0.01g FeCl_3_·6H_2_O, 15.0g agar.

**Sphaerotilus Defined medium** (Atlas 2010)

Sphaerotilus Defined medium (L^-1^ distilled water): 15.0g agar, 5.0g glycerol, 0.9g glutamic acid, 0.5g FeSO_4_·7H_2_O, 0.1g MgSO_4_·7H_2_O, 0.03g CaCl_2_·2H_2_O, 0.03g ZnSO_4_·7H_2_O, 100.0mL phosphate solution (5.7g K_2_HPO_4_, 2.3g KH_2_PO_4_ per 500.0mL).

**Synthetic Sea Water medium 1** (Yakimov et al., 1998)

SM I contained (L^-1^ distilled water): 10 g pyruvate, 23g NaCl, 0·75g KCl, 1.47g CaCl_2_.2H_2_O, 5·08g MgCl_2_.6H_2_O, 6.16g MgSO_4_.7H_2_O, 0·89g Na_2_HPO_4_.2H_2_O, 5.0g NaNO_3_, and 0·03g FeSO_4_.7H_2_O.

**TCBS medium** (Atlas 2010)

TCBS medium contained (L^-1^ distilled water): 20.0g sucrose, 10.0g NaCl, 10.0g sodium citrate, 10.0g Na_2_S_2_O_3_, 5.0g yeast extract, 5.0g pancreatic digest of casein, 5.0g peptic digest of animal tissue, 5.0g oxgall, 3.0g sodium cholate, 1.0g ferric citrate, 0.04g thymol Blue, 0.04g bromthymol blue, 14.0g agar.

**XLD Agar** (Atlas 2010)

XLD Agar contained (L^-1^ distilled water):7.5g lactose, 7.5g sucrose, 6.8g Na_2_S_2_O_3_, 5.0g L-lysine, 5.0g NaCl, 3.5g xylose, 3.0g yeast extract, 2.5g sodium desoxycholate, 0.8g ferric ammonium citrate, 0.08g phenol red, 13.5g agar.

**Extraction of metagenomes and metatranscriptome**

Immediately after the samples collected in an exploration reached the laboratory, each 0.22 μm mixed cellulose ester membrane, which was used on-field to filter out microbial cells from 1 L vent-water and was subsequently hydrated and preserved in 5 mL of 50 mM Tris:EDTA (TE pH 7.8) for the purpose of metagenome extraction, was shredded with sterile scissors inside the 10 mL cryovial in which it was brought from the field (Mondal et al., 2024). All the 10 vials lined up in this way in relation to a given sampling occasion were vortexed for 15 minutes, and the filter-shreds were allowed to settle down at the bottom of the vials. The supernatants were then collected in a fresh sterilized vial without disturbing the debris - total volume of cell suspension pooled in this way from the 10 individual vials was approximately 50 mL. This 50 mL cell suspension was centrifuged for 30 min at 10,800 *g*, subsequent to which the upper 48 mL TE solution was discarded. The bottom 2 mL - apparently containing all the microbial cells that were there in the 10 L vent-water filtered on the sampling occasion - was subjected to metagenomic DNA extraction using PureLink Genomic DNA Mini Kit (Thermo Fisher Scientific, USA). Over the six different metagenome extraction procedures carried out in this way, 30-100 ng DNA could be finally eluted in 30 µL nuclease-free water, with each preparation corresponding to the total environmental DNA present in 10 L CMSI vent-water. In tandem with each metagenome extraction procedure, purported kitome DNA was prepared using the same PureLink kit without adding any CMSI cell suspension, instead of which 2 mL sterilized nuclease-free water was used. Micro-volume spectroscopy using neither NanoDrop nor Qubit technology (both from Thermo Fisher Scientific, USA) detected any DNA in the final eluent of any of the so-called kitomes prepared; therefore, none of the six kitome eluents was subjected to sequencing.

In order to obtain total RNA from the microbial cells that were still adhered to the mixed cellulose ester membranes which were used on-field to filter out cells from batches of 1 L vent-water, the membranes were inserted into Power Water DNA Bead Tubes provided with RNeasy Power Water Kit (Qiagen GmbH, Germany), and total RNA was extracted following the manufacturer’s protocol. To extract total RNA from the microbial cells that had already got dislodged from the membranes and were floating in RNAlater, the solutions contained in the cryovials brought from the field were pooled to get an approximately 250 mL potential cell suspension. This cell suspension was passed through a fresh 0.22 μm mixed cellulose ester membrane, and the membrane with its cellular residue was inserted into Power Water DNA Bead Tubes to extract total RNA as mentioned above. The two separate RNA preparations were pooled, concentrated, and finally resuspended in 30 µL RNase-free water, yielding a final RNA concentration of 4.4 ng µL^-1^. RNA integrity number (RIN) was assayed for this final stock solution using a 2100 Bioanalyzer System (Agilent Technologies, USA), and found to be 7.1. In tandem with the above procedure, purported kitome RNA was prepared using the same RNeasy kit, RNAlater reagent, and 0.22 μm mixed cellulose ester membrane, albeit without filtering any CMSI vent-water or cell suspension through the membrane. Micro-volume spectroscopy using neither NanoDrop, nor Qubit or Bioanalyzer, technology detected any RNA in the final eluent of the so-called kitome prepared. The kitome eluent, therefore, was not sequenced subsequently.

**Supplementary Results and Discussion**

**Molecular drivers underlying the asymmetric ecology of putative hyperthermophiles**

The overwhelming numeral as well as functional predominance of Aquificia_2 coincided with its expression of a high number (171) of metabolic modules (Figure 10), and possession of the highest EL_cc_ values for 16 out of the total 23 COG-categories defined within the 66 CMSI MAGs/genomes (Figure 8). In particular, the EL_cc_ values recorded for Energy production and conversion; and Secondary metabolites biosynthesis, transport and catabolism, in Aquificia_2 were far higher than (i) the EL_cc_ of these two COG-categories in other CMSI species, as well as (ii) the EL_cc_ of all other COG-categories within Aquificia_2. The Thermoproteales species - which slightly lagged Aquificia_2 in terms of Φ but trailed more than four times in terms of Π_μ_ - possessed the next highest EL_cc_ for Energy production and conversion (Figure 8). Species having sequentially lower Φ values had commensurately lower EL_cc_ for Energy production and conversion, irrespective of the magnitude of their Π_μ_. Concurrently, across the 66 CMSI species, EL_CC_ for Energy production and conversion exhibited a higher level of positive correlation with Φ (*R*^2^ = 0.84; *P* = < 0.001), compared to Π_μ_ (*R*^2^ = 0.7; *P* = < 0.001). Φ values had *R*^2^ > 0.9, alongside *P* = 0, with the EL_CC_ values of all COG-categories except Cell wall/membrane/envelope biogenesis; Defense mechanisms; RNA processing and modification; and Transcription (*R*^2^ was 0.4-0.84, and *P* value was 0, for the last four categories). Counter to these, COG-categories influencing Π_μ_ were few. EL_CC_ values recorded for Cell wall/membrane/envelope biogenesis; Intracellular trafficking, secretion, and vesicular transport; and Energy production and conversion, showed maximum positive correlations with Π_μ_ (*R*^2^ = 0.73, 0.72, and 0.7, alongside *P* = 0, respectively). EL_CC_ values of COG-categories except these three showed *R*^2^ < 0.7, alongside *P* = 0, with Π_μ_ (Table S79).

The Thermoproteales species exhibited *in situ* functionality as high as Aquificia_2, even though its prevalence over the same sampling period was approximately one-fourth that of Aquificia_2. Thus, functionality:prevalence (Φ:Π_μ_) ratio, a measure of overall ecological efficacy of a species, was >1.2 for Aquificia_2 but 4.3 for the Thermoproteales member (Figures 5C and S2). The extraordinary ecological efficacy of the latter coincided with its exceptionally higher EL_cc_ for RNA processing and modification, and Extracellular structures, compared to all other CMSI species (relevant genes were mostly absent in the Aquificia MAGs; Figure 8). The Thermoproteales species also exhibited higher EL_cc_ for Cell cycle control, cell division, chromosome partitioning; Defense mechanisms; Mobilome: prophages, transposons; Signal transduction mechanisms; and Transcription, compared to not only Aquificia_2 but also every other CMSI species. Furthermore, ELcc for Energy production and conversion; and Secondary metabolites biosynthesis, transport and catabolism, was much higher in the Thermoproteales species, compared to all other CMSI inhabitants except Aquificia_2. Within the expressed gene-catalog of this Thermoproteales member, the CDS for alkyl hydroperoxide reductase (AhpC, which also acts as a protein chaperone under stress; Chuang et al., 2006) had the highest EL_gene_ (Table S11). Homologs of *ahpC* being either absent or having very low EL_gene_ in the other putative hyperthermophiles of the habitat, the Thermoproteales species was apparently better protected from the oxidative stressors typical of geothermal ecosystems (Imlay 2013).

Thermoprotei_1A and Thermoprotei_1B had Φ values five to six times lower than those of Aquificia_2 or Thermoproteales. However, these functionality levels corresponded to high Π_μ_ in Thermoprotei_1A and low Π_μ_ in Thermoprotei_1B, so Φ:Π_μ_ for the two archaea were 0.3 and 1.1 respectively (Figures 5C and S2). The relatively higher ecological efficacy of Thermoprotei_1B, compared to Thermoprotei_1A, could among other qualities be attributable to the former’s greater efficiency in genome maintenance and stress management. These aptitudes of Thermoprotei_1B are highlighted by its exceedingly high EL_gene_ for *tatD* (compare Tables S12 and S10; TatD mediates cleanup of DNA breaks and errors; Chen et al., 2014) and exclusive metatranscriptomic signatures for the biosynthesis and/or metabolism of phosphoribosyl pyrophosphate, proline, UDP-2,3-diacetamido-2,3-dideoxy-α-D-mannuronic acid, and uridine diphosphate-N-acetylgalactosamine (Figure 10), which are known to be associated with different stress mitigation mechanisms (Kawamura et al., 1985; Zhang et al., 2015; Hove-Jensen et al., 2016). In contrast, Thermprotei_1A, despite its higher prevalence, could not render an overall functionality above the level of Thermprotei_1B apparently due to, among other reasons, lower EL_gene_ for *rpoZ* (RNA polymerase subunit K/omega; Gunnelius et al., 2014) and *spt4* (transcription elongation factor; Huffines et al., 2021). However, exceedingly high *livK*-mediated peptides and amino acids sequestration (compare Tables S10 and S12; Ribardo and Hendrixson 2011) and Intracellular trafficking, secretion, and vesicular transport (Figure 8) might have allowed Thermprotei_1A to maintain a prevalence-level disproportionately higher than its functionality.

Aquificia_1 and Aquificia_3 had equivalently lower functionality and prevalence, compared to Aquificia_2; yet, they were the most abundant and active inhabitants of the vent after the four CCPMS-I members. Aquificia_1 and Aquificia_3, despite expressing equivalent number of KEGG Modules as Aquificia_2 (Figure 10), lagged in terms of prevalence as well as functionality apparently due to their low EL_cc_ for Cell cycle control, cell division, chromosome partitioning; Replication, recombination and repair; and Transcription (Figure 8).

The remaining eight putative hyperthermophiles of CMSI had Φ < 30 and Π_μ_ < 50, so unequivocally lagged far behind the six species discussed above in terms of functionality as well as prevalence. That said, Thermoprotei_2 and *Pyrobaculum*_*aerophilum*, by virtue of certain transcriptomic specialties, had Φ:Π_μ_ ratios >1 (Figures 5C and S2). Molecular underpinnings of the numeral/functional laggardness of the ecologically less-efficient archaea are discussed below, in conjunction with whatever adaptational advantages they had, and molecular strategies they employed, for their sustenance in the CMSI vent-water.

**Molecular underpinnings of the numeral/functional laggardness of the ecologically less-efficient archaea**

Among the eight archaea having Φ <30 alongside Π_μ_ <50 (Table S8; Figure 5C), only Thermoprotei_2 and *Pyrobaculum*_*aerophilum* had Φ:Π_μ_ ratios >1 (1.6 and 1.7 respectively; Figure 2).

Of all the CDSs expressed by Thermoprotei_2, the ones encoding the methyl-accepting chemotaxis protein Tar, and the oxidative-damage-protecting DNA-binding protein Dps, had the highest EL_gene_ values. Concurrently, CDSs of the transporters DppB, DppC, and PstS also had high EL_gene_ values (Table S20). It, therefore, seemed likely that *in situ* adaptations of this archaeon include quest for genome stability (Pulliainen et al., 2005; Tatur et al., 2007), and acquisition of relevant inorganic nutrients (Pletzer et al., 2014; Ranjit et al., 2024); plus chemotaxis along gradients of nutrients, redox substrates, temperature, and/or other biophysical factors (Cha et al., 2022). Comparative laggardness of Thermoprotei_2 in terms of prevalence concurred with its low EL_cc_ for Cell cycle control, cell division, chromosome partitioning; and Replication, recombination and repair. Simultaneously, its overall low expressivity across metabolisms corresponded to the low EL_cc_ for Transcription (Table S75; Figure 8). Furthermore, genes for hydrogenase were missing from the MAG, while the modules present for cytochrome oxidase, and dissimilatory reductions of nitrate to ammonium, and elemental sulfur to sulfide, were not expressed *in situ*. Towards ATP generation, only the genes concerned with peptides-/amino-acids-based substrate-level phosphorylation with the help of aldehyde:ferredoxin oxidoreductase were expressed at low levels - apparently, exclusive reliance on this low-efficiency pathway of energy transduction (Mayer and Müller, 2014) was a key reason behind the comparatively lower activity and prevalence of Thermoprotei_2.

Among all the CMSI archaea, *Pyrobaculum*_*aerophilum* exhibited comprehensive metatranscriptomic signatures for the highest number (163) of KEGG Modules (Table S76), yet its functionality as well as prevalence was lower, compared to the Thermoproteales species, Thermoprotei_1A, Thermoprotei_1B, and Thermoprotei_2. Within the expressed CDS-catalog of this archaeon, highest EL_gene_ was recorded for Cob(II)alamin adenosyltransferase. Since this enzyme is critical for the activation of cofactors required for carbon-skeleton rearrangement (Mera and Escalante-Semerena, 2010), its high level of expression (Table S25) suggested that the CMSI vent-water is not the habitat of choice for *Pyrobaculum*­­_*aerophilum* and the archaeon is facing adaptive stress *in situ*. As a result, the organism is spreading its biomacromolecular resources thinly across diverse metabolic processes. Corroborative to this hypothesis, *Pyrobaculum*_*aerophilum* expressed the bifunctional guanosine tetraphosphate/pentaphosphate [(p)ppGpp] synthase/hydrolase SpoT (no *spoT* homolog was detected in any of the other 10 CMSI archaea). This, in turn, showed that *Pyrobaculum*_*aerophilum* was rendering such stress-induced stringent responses *in situ* that involved RNA polymerase reconfiguration towards enhanced transcription of survival-related genes, rather than those promoting growth (Imlay 2003; Magnusson et al., 2005; Jang and Imlay, 2007).

The six archaeal species identified as Thermoprotei_3, Thermoprotei_4, Thermoproteota_1, Thermoproteota_2, Thermoproteota_3, and Thermoproteota_4 had Φ <6 alongside Π_μ_ <15 (Table S8; Figure 5C). Modules for guanine ribonucleotide degradation and non-phosphorylative Entner-Doudoroff pathway were missing in these six archaeal MAGs, whereas the archaea having Φ >6 plus Π_μ_ >15 exhibited comprehensive metatranscriptomic signatures for these two pathways. Inability to convert GMP to urate, or gluconate/galactonate to glycerate, might lead to the accumulation of oxidized guanine nucleotides under high-temperature stress (Sekiguchi et al., 2013, Singh et al., 2019). Absence of non-phosphorylative Entner-Doudoroff pathway can further reduce the ability to catabolize glucose/gluconate and produce ATP (Reher et al., 2010), thereby rendering these archaea ecologically uncompetitive. Furthermore, these six archaea, except Thermoprotei_3 and Thermoproteota_2, either lacked pyruvate oxidation or did not express the module (Table S76). Consequently, they might be facing a persistent scarcity of acetyl-CoA for energy supply and biomacromolecule synthesis (Ma et al., 1997; Richter et al., 2024), especially since none of them except Thermoprotei_3 possessed or expressed the pentose phosphate pathway, or for that matter the alternative ribulose monophosphate pathway. Shortage of ribose-5-phosphate can also jeopardise different reductive biosynthesis processes, nucleotide synthesis, carbon homeostasis, and oxidative stress management (Orita et al., 2006; Stincone et al., 2015). Thermoprotei_3, apart from expressing pyruvate oxidation, pentose phosphate pathway, and alternative ribulose monophosphate pathway *in situ*, exhibited comprehensive metatranscriptomic signature for the glycine cleavage module (Table S76), and exceedingly high EL_gene_ for 5,10-methylenetetrahydrofolate reductase (Table S23). Collectively, these metabolic capacities can enhance biomass generation, which in turn can lead to higher prevalence *in situ*.

**Strategies underlying the ecological fitness of the putative thermophiles and potential moderate-thermophiles** **of CMSI**

Metabolic machineries of Thermaceae_1, Thermaceae_2, and the two species ascribed to *Thermoflexus* and Thermoflexaceae (Tables S16, S15, S28, and S21, respectively), seemed to be focused on DNA, RNA and/or protein quality control, because they possessed at least four of the following chaperone-coding genes among their CDSs with highest expression-levels: *clpA*, *cspC*, *dnaK*, *dnaJ*, *ibpA*, *groEL*, *groES*, *truB*, and *yugP*.

Among the 27 CMSI species affiliated to taxa having few moderately-thermophilic members, the two entities having highest Φ and Π_μ_, namely Armatimonadota_1 and the *Thermosynechococcus* species, had *dnaK*, *groEL* and/or *groES*, plus the oxidative-stress-mitigating *ahpC*, *ahpF*, *dps* and/or *yrkE*, among their CDSs having highest EL_gene_ (Tables S17 and S19). The other potential moderate-thermophiles detected also encompassed a number of chaperones and/or oxidative-stress-mitigating genes among their CDSs with highest expression-levels.

Compared to the putative thermophiles and hyperthermophiles that lagged behind them in terms of Φ as well as Π_μ_, Thermaceae_1 and Thermaceae_2 had higher EL_CC_ for three and 15 COG-categories respectively (Figure 8). Thermaceae_1 and Thermaceae_2 also exhibited comprehensive metatranscriptomic signatures for the modules rendering 2-oxocarboxylic acid chain extension; formaldehyde assimilation; sulfur oxidation; and biosynthesis of coenzyme M, lysine, pyocyanine, and tetrahydrofolate (Figure 10); these modules were either missing in the five hyperthermophiles and two thermophiles lagging Thermaceae_1 and Thermaceae_2 in terms of Φ as well as Π_μ_ (Figure 5C), or were not expressed by them. Most of these modules could contribute to redox homeostasis, energy metabolism, and detoxification of reactive intermediates under physicochemically unfavorable conditions. Furthermore, Thermaceae_1 and Thermaceae_2 (Table S15 and S16), but not the hyperthermophiles (Tables S22, S29, S34, S35, and S38) and thermophiles (Tables S21 and S28) lagging them in terms of Φ and Πμ, had the following genes concerned with DNA and protein quality control among their CDSs with highest EL_gene_ values: tRNA U55 pseudouridine synthase maintaining translation fidelity under stress *truB* (Keffer-Wilkes et al., 2016), small heat shock protein of the HSP20 family *ibpA* (Miwa and Taguchi, 2023), single-stranded DNA-binding protein *ssb*, HSP70 molecular chaperone *dnaK*, DnaJ-class co-chaperone *dnaJ* (Susin et al., 2006), ATP-dependent Clp protease subunit *clpA* (Capestany et al., 2008), and cold shock protein of the CspA family *cspC* (Keto-Timonen et al., 2016).

The *Thermoflexus* and Thermoflexaceae species, together with Armatimonadota_1, exhibited comprehensive metatranscriptomic signatures for the modules C4-dicarboxylic acid cycle; formaldehyde assimilation; biosynthesis of fatty acid, galactofuranan, glutathione, triacylglycerol; and degradation of guanine ribonucleotide and sphingosine (Figure 10). These modules were either missing in the five putative hyperthermophiles lagging these three bacteria in terms of Φ (Figure 5C) or were not expressed by them. Several of these modules are known to mitigate stress by rendering antioxidant functions, neutralizing reactive intermediates, and adjusting redox balance, within microbial cells (Forman et al., 2009). Furthermore, the *Thermoflexus* (Table S21) and Thermoflexaceae (Table S28) species, together with Armatimonadota_1 (Table S17) - but not the putative hyperthermophiles lagging these three bacteria in terms of Φ (Tables S22, S29, S34, S35, and S38) - had the chaperones *clpA*, *dnaK*, and/or *ibpA* at the top of their *in-situ*-transcribing CDS-catalogs in terms of EL_gene_ values.

The *Thermosynechococcus* species had higher EL_CC_ for 11 COG-categories, compared to the 11 putatively hyperthermophilic and thermophilic species that lagged behind it in terms of Φ as well as Π_μ_ (Figure 8). Apart from the genes for the molecular chaperones DnaK, GroEL, and/or GroES, and the oxidative stress management proteins AhpC, AhpF, Dps and/or YrkE, the CMSI *Thermosynechococcus* (Table S19) exhibited comprehensive metatranscriptomic signatures for the KEGG Modules rendering sulfate-sulfur assimilation, and biosynthesis of ADP-LDmanHep, biotin, GDP-DDmanHep, lactosylceramide, KDO2-lipid A, O-glycan, pyridoxal phosphate, tocopherol, or tocotorienol, and zeaxanthin (Figure 10). These modules were either missing from the genomes of the seven hyperthermophiles (Tables S22, S23, S25, S29, S34, S35, and S38) and four thermophiles (Tables S15, S16, S21, and S28,) lagging the *Thermosynechococcus* species in terms of Φ as well as Π_μ_ (Figure 5C), or were not expressed by them. Many of these modules are known to stabilize membrane‑associated proteins, mitigate permeability stress, and sustain essential signalling processes under adverse physicochemical conditions (Grabowicz and Silhavy, 2017, Willdigg and Helmann, 2021). Furthermore, the *Thermosynechococcus* species (Table S19), but not the 11 laggard hyperthermophiles and thermophiles, had *ahpC*, *dps*, cytochrome *b* subunit *qcrB*/*petB*, and cytochrome *c*-550 *cccA*, among its CDSs with highest EL_gene_ values.

**Molecular drivers of the persistence exhibited by the CMSI mesophiles**

Of the 17 mesophiles detected in the CMSI vent-water, 13 invariably expressed at least some genes concerned with Energy production and conversion; Transcription; Translation, ribosomal structure and biogenesis; and Posttranslational modification, protein turnover, chaperones, apart from genes related to other COG-categories (Figure 8). No gene was expressed under the COG-category Cell cycle control, cell division, chromosome partitioning by any CMSI mesophile other than *Stutzerimonas* CFSM4_SD3 (Table S36), *Micrococcus* LASM2_SD2 (Table S50), and *Mycolicibacterium* BG11O_SDJG (Table S64), which in turn expressed only very few genes under this category (*Stutzerimonas* expressed five genes, including *ftsZ*, but not *smc*, *ftsN*, or *cdvB*; *Micrococcus* and *Mycolicibacterium* expressed only *rodZ* and *ftsK* respectively; Figure 9). Concurrent with the above data, the *Stutzerimonas* and *Mycolicibacterium* isolates were the only CMSI mesophiles for which metatranscriptomic signatures of defined KEGG Modules were recorded (Figure 10); and Φ values of the *Stutzerimonas* and *Micrococcus* isolates (plus that of *Acinetobacter_indicus*) were orders of magnitude higher than the Φ recorded for all the other mesophiles (Figure 5C).

Among the CMSI mesophiles, *Stutzerimonas* CFSM4_SD3 had the highest EL_CC_ for nine out of the total 19 COG-categories detected across these 17 species (Figure 8). The *Stutzerimonas* isolate exhibited metatranscriptomic signatures for total 82 modules (Figure 10), of which 29 were associated with adaptation to thermal, oxidative, nutritional, and other stressors. Examples of these modules included β‑oxidation of fatty acids; biosynthesis of glycogen, polyamine, and trehalose; and dimethylsulfoniopropionate degradation, all of which are well appreciated for their involvement in energy conservation, osmoprotection, proteins/membranes stabilization, and other biophysical strategies of stress management (Kaushik et al., 2003; Fujita et al., 2007; Oshima 2007; Wilson et al., 2010; Wang et al., 2022).

*Mycolicibacterium* BG11O_SDJG exhibited metatranscriptomic signatures for 17 modules (Figure 10), of which 11 had direct or indirect roles in thermal/oxidative stress management. Examples included β‑oxidation of fatty acids; biosynthesis of fatty acid, and trehalose; dimethylsulfoniopropionate degradation; inositol phosphate metabolism (Kaushik et al., 2003; Fujita et al., 2007; Wang et al., 2009; Belfquih et al., 2022; Fuangthong and Helmann 2002).

Among the 17 CMSI mesophiles, *Micrococcus* LASM2_SD2 had the highest EL_CC_ for Coenzyme transport and metabolism; Inorganic ion transport and metabolism; and Defense mechanism. Individually, this bacterium had high EL_gene_ for OsmC/OhrA peroxidase, nitroimidazole reductase NimA, DJ-1 deglycase, and a DNA methylase of Type I restriction-modification system (Table S50), implicating *in situ* mitigation of stress-induced damage to biomacromolecules (Leiros et al., 2004; Mihoub et al., 2017; Zhu et al., 2022).

Among the 17 mesophiles, *Acinetobacter_indicus* had the highest EL_CC_ for Energy production and conversion; Mobilome: prophages, transposons; Nucleotide transport and metabolism; Secondary metabolites biosynthesis, transport and catabolism; Transcription; and Translation, ribosomal structure and biogenesis (Figure 8). This bacterium had high EL_gene_ for the polyhydroxyalkanoate-granule-associated phasin PhaF (Table S54), indicating protection against extreme temperature, oxidative stress, and osmotic flux (Tarazona et al., 2020; Obulisamy et al., 2021).

**Supplementary Table**

**Table S3.** Species-level entities discovered in the form of metagenome-assembled genomes (MAGs), over the six different explorations of the CMSI vent-water; maximum temperature (Tmax °C) known for the laboratory growth of the member strains of the lowest-possible taxon to which every MAG-based species could be classified has also been given.

| **Identification of the species** | **Metagenomic datasets^§^ from where sibling MAGs could be reconstructed** | | | | | | **Maximum temperature (T_max_) known for laboratory growth of any member of the taxon up to which the species could be classified** | | |
| --- | --- | --- | --- | --- | --- | --- | --- | --- | --- |
|  | **1** | **2** | **3** | **4** | **5** | **6** | **T_max_ (°C)** | **Species where the T_max_ was recorded** | **Reference** |
| MAG_*Pyrobaculum*_*aerophilum* |  |  | + | + | + | + | 104 | *Pyrobaculum aerophilum* | Huber et al. (1993*)* |
| MAG_Thermoproteales | + | + | + | + | + | + | 104 | *Pyrobaculum aerophilum* | Huber et al. (1993) |
| MAG_Thermoprotei_1A | + | + |  | + | + | + | 121 | *Geogemma barossii* | Kashefi and Lovely (2004) |
| MAG_Thermoprotei_1B | + | + |  | + | + | + |  |  |  |
| MAG_Thermoprotei_2 | + | + | + | + | + | + |  |  |  |
| MAG_Thermoprotei_3 |  |  | + | + | + | + |  |  |  |
| MAG_Thermoprotei_4 |  |  |  |  | + |  |  |  |  |
| MAG_Thermoproteota_1 | + | + | + | + | + | + | 121 | *Geogemma* *barossii* | Kashefi and Lovely (2004) |
| MAG_Thermoproteota_2 |  |  | + | + |  |  |  |  |  |
| MAG_Thermoproteota_3 |  |  |  | + |  |  |  |  |  |
| MAG_Thermoproteota_4 |  |  |  |  | + | + |  |  |  |
| MAG_Aquificia_1 | + | + | + | + | + | + | 95 | *Aquifex pyrophilus* | Huber et al. (1992) |
| MAG_Aquificia_2 | + | + | + | + | + | + |  |  |  |
| MAG_Aquificia_3 | + | + | + | + | + | + |  |  |  |
| MAG_Thermaceae_1 |  | + |  | + | + |  | 85 | *Thermus thermophilus* | Oshima et al. (1974) |
| MAG_Thermaceae_2 |  |  |  |  |  | + |  |  |  |
| MAG_Thermoflexaceae | + | + | + | + | + |  | 75 | *Thermoflexus hugenholtzii* | Dodsworth et al. (2014) |
| MAG_*Thermoflexus* |  | + | + | + |  | + | 75 | *Thermoflexus hugenholtzii* | Dodsworth et al. (2014) |
| MAG_Cyanobacteriota_1 |  |  |  |  | + |  | 72 | *Thermostichus lividus* | Meeks and Castenholz (1971) |
| MAG_Cyanobacteriota_2 |  |  |  |  | + |  |  |  |  |
| MAG_Armatimonadota_1 | + | + | + | + | + | + | 73 | *Chthonomonas calidirosea* | Lee et al. (2011) |
| MAG_Armatimonadota_2 |  | + |  | + | + | + |  |  |  |
| MAG_Gammaproteobacteria |  |  |  |  | + |  | 68 | *Inmirania thermothiophila* | Slobodkina et al. (2016) |
| MAG_*Thermosynechococcus* | + | + | + | + | + | + | 60 | *Thermosynechococcus elongatus* | Onai et al. (2004) |
| MAG_*Georgenia* |  |  |  |  | + | + | 60 | *Georgenia sediminis* | You et al. (2013) |
| MAG_*Microbacterium* * |  |  | + | + |  |  | 50 | *Microbacterium sediminis* | Yu et al. (2013) |
| MAG_*Brevundimonas*_1 |  |  |  |  | + |  | 50 | *Brevundimonas naejangsanensis* | Kang et al. (2009) |
| MAG_*Brevundimonas*_2 * |  |  |  |  |  | + |  |  |  |
| MAG_*Brevundimonas*_3 |  |  |  |  |  | + |  |  |  |
| MAG_Burkholderiales_1 |  |  | + |  |  |  | 65 | *Thiomonas thermosulfata* | Shooner et al. (1996) |
| MAG_Burkholderiales_2 |  |  | + |  |  |  |  |  |  |
| MAG_Burkholderiales_3 |  |  | + |  |  |  |  |  |  |
| MAG_Halomonadaceae | + | + | + | + | + | + | 60 | *Halomonas zhaodongensis* | Jiang et al. (2013) |
| MAG_Sphingomonadales |  |  |  |  | + |  | 60 | *Novosphingobium endophyticum* | Li et al. (2015) |
| MAG_*Tepidimonas*_*ignava* | + |  |  |  |  |  | 64 | *Tepidimonas ignava* | Moreira et al. (2000) |
| MAG_*Vibrio*_*metschnikovii* | + | + |  | + |  | + | 50 | *Vibrio gangliei* | Meng et al. (2018) |
| MAG_Moraxellaceae |  |  | + |  |  |  | 45 | *Acinetobacter larvae* | Liu et al. (2017) |
| MAG_*Psychrobacter* |  |  | + |  |  |  | 40 | *Psychrobacter celer* | Yoon et al. (2005) |
| MAG_*Acinetobacter*_*indicus* |  | + |  |  |  |  | 45 | *Acinetobacter larvae* | Liu et al. (2017) |
| MAG_*Agrobacterium*_*pusense* * |  |  |  |  | + | + | 42 | *Agrobacterium pusense* | Castellano-Hinojosa et al. (2021) |
| MAG_*Sphingomonas* |  |  |  |  | + | + | 45 | *Sphingomonas zeicaulis* | Gao et al. (2016) |
| MAG_Boseaceae |  |  |  |  |  | + | 42 | *Bosea minatitlanensis* | Ouattara et al. (2003) |
| MAG_Bacteria_1 | + | + | + | + | + | + | NA | NA | NA |
| MAG_Bacteria_2 |  |  | + |  |  |  | NA | NA | NA |
| MAG_Bacteria_3 |  |  | + | + |  |  | NA | NA | NA |
| MAG_Bacteria_4 |  |  |  | + |  |  | NA | NA | NA |

^§^	Metagenomic datasets indicated as 1 through 6 represent FN_Nov_21_Mg, AN_Nov_21_Mg, FN_Apr_22_Mg, AN_Apr_22_Mg, FN_Nov_22_Mg, and AN_Nov_22_Mg, respectively.

*	Corresponding to these three MAGs, genomically indistinguishable strains (having 97.2-100% OrthoANI and 74.7-99.7% dDDH) were also isolated from the CMSI vent-water sample collected through 01-Nov-2022 (see Figure 3).

**Supplementary Figures**

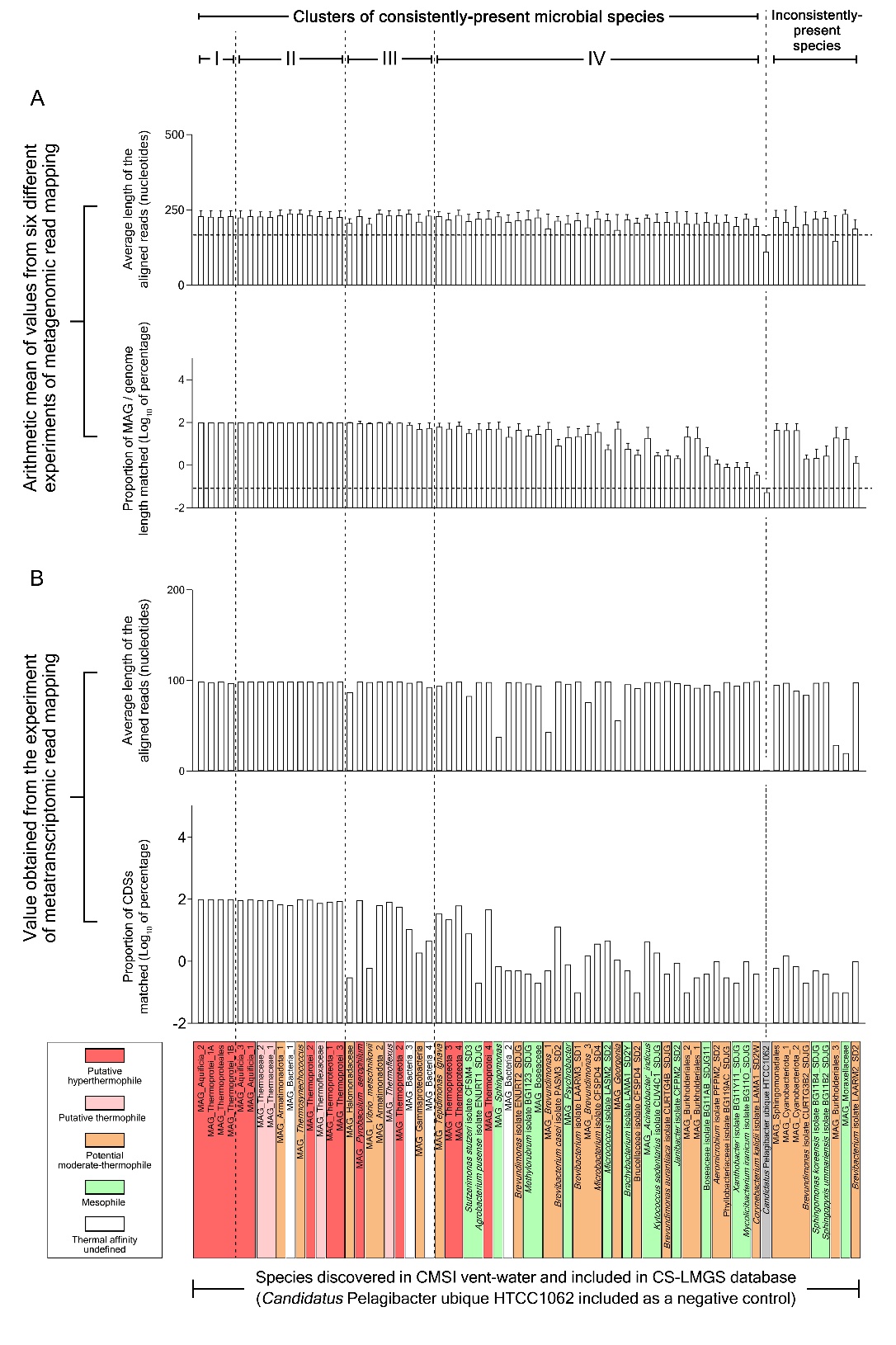

**Figure S1.** Alignment statistics underlying the results obtained for the metagenomic and metatranscriptomic read mapping experiments conducted upon the CS-LMGS database. For each species included in the CS-LMGS database, (**A**) plots the arithmetic mean of the average lengths of the aligned reads (nucleotides) recorded over the six different experiments of metagenomic read mapping; (**B**) plots the arithmetic mean of the percentages of its MAG/genome length that were matched over the six different experiments of metagenomic read mapping. In panels “A” and “B”, error bars indicate the standard deviations of the data. For each species included in the CS-LMGS database, (**C**) plots the average length of the aligned reads (nucleotides) recorded in the metatranscriptomic read mapping experiment; (**D**) plots the percentage of its annotated CDSs that was matched in the metatranscriptomic read mapping experiment. Species names are highlighted by different colors based on their putative thermal affinities (the color code is same as the one used in Figures 5 through 10).

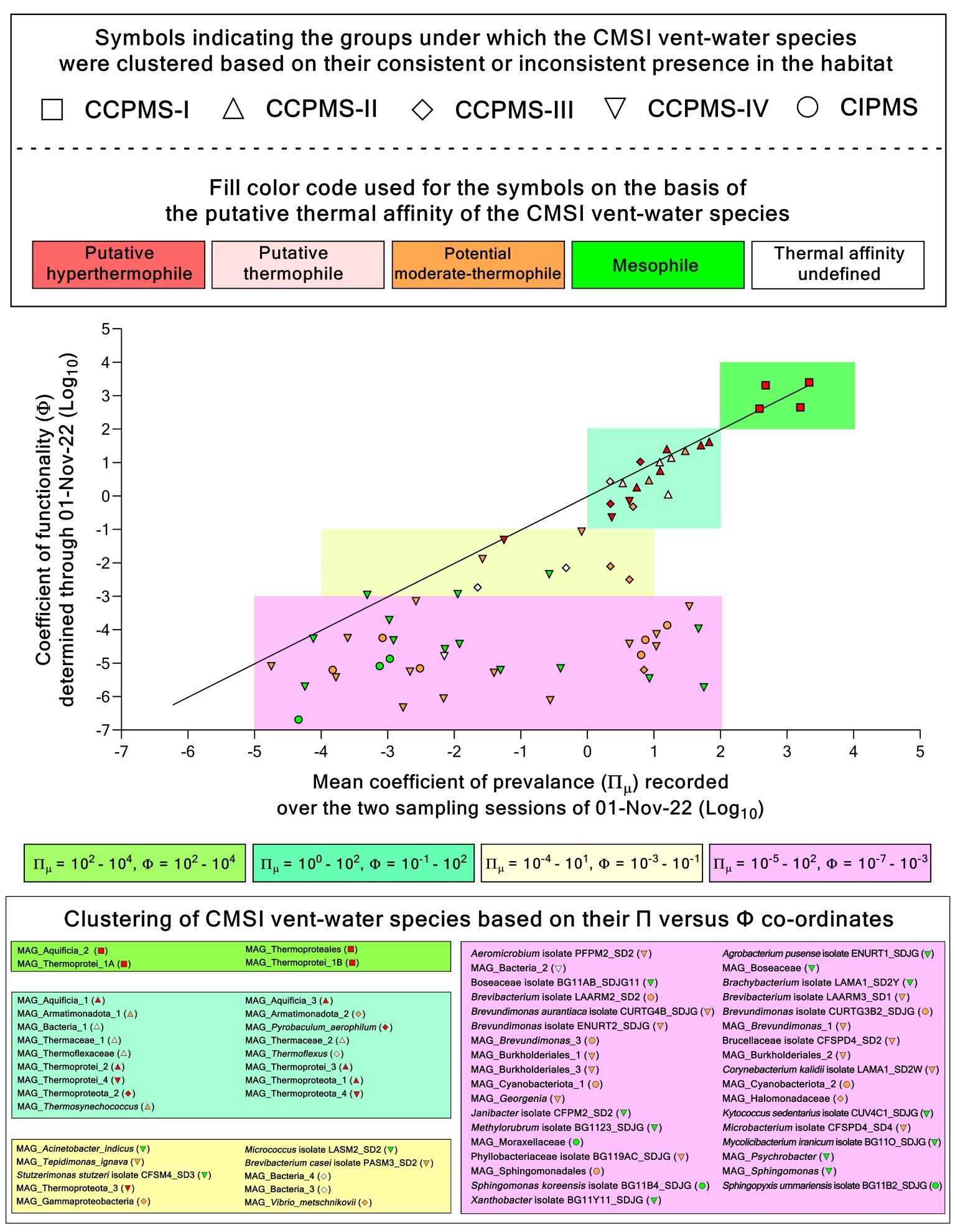

**Figure S2.** Scatter plot showing the Φ of each CMSI species as a function of its Π_μ_. Specific regions of the plot, defined by distinct ranges of Φ and Π_μ_ values, are demarcated by different shades to encompass species having similar Φ:Π_μ_ ratios. A linear regression line, constrained to pass through the origin, is overlaid on the plot to delineate the points having Φ:Π_μ_ = 1.
